# Kinetic-model-guided engineering of multiple *S. cerevisiae* strains improves *p*-coumaric acid production

**DOI:** 10.1101/2024.12.15.628543

**Authors:** Bharath Narayanan, Wei Jiang, Shengbao Wang, Javier Sáez-Sáez, Daniel Weilandt, Maria Masid Barcon, Viktor Hesselberg-Thomsen, Irina Borodina, Vassily Hatzimanikatis, Ljubisa Miskovic

## Abstract

The use of kinetic models of metabolism in design-build-learn-test cycles is limited despite their potential to guide and accelerate the optimization of cell factories. This is primarily due to difficulties in constructing kinetic models capable of capturing the complexities of the fermentation conditions. Building on recent advances in kinetic-model-based strain design, we present the rational metabolic engineering of an *S. cerevisiae* strain designed to overproduce *p*-coumaric acid (*p*-CA), an aromatic amino acid with valuable nutritional and therapeutic applications. To this end, we built nine kinetic models of an already engineered *p*-CA-producing strain by integrating different types of omics data and imposing physiological constraints pertinent to the strain. These nine models contained 268 mass balances involved in 303 reactions across four compartments and could reproduce the dynamic characteristics of the strain in batch fermentation simulations. We used constraint-based metabolic control analysis to generate combinatorial designs of 3 enzyme manipulations that could increase p-CA yield on glucose while ensuring that the resulting engineering strains did not deviate far from the reference phenotype. Among 39 unique designs, 10 proved robust across the phenotypic uncertainty of the models and could reliably increase *p*-CA yield in nonlinear simulations. We implemented these top 10 designs in a batch fermentation setting using a promoter-swapping strategy for down-regulations and plasmids for up-regulations. Eight out of the ten designs produced higher *p*-CA titers than the reference strain, with 19 – 32% increases at the end of fermentation. All eight designs also maintained at least 90% of the reference strain’s growth rate, indicating the critical role of the phenotypic constraint. The high experimental success of our in-silico predictions lays the foundation for accelerated design-build-test-learn cycles enabled by large-scale kinetic modeling.

## 1 Introduction

Systems metabolic engineering enables us to rewire microorganisms to produce valuable chemicals, such as biofuels, aromatic amino acids, and other plant derivatives, in an environmentally friendly manner^1–3^. A common challenge in establishing microbial cell factories is optimizing the strain to ensure cost-effective production. There are two broad approaches to strain optimization: evolutionary and rational. Evolutionary approaches such as adaptive laboratory evolution^4^ (ALE) and random mutagenesis^5^ use the power of natural selection to choose overproducing strains. Conversely, rational strain design targets enzymes deliberately chosen using existing knowledge of metabolic and regulatory networks, as well as computational tools^6^. Common rational strategies include the removal of inhibitory feedback loops, downregulating genes involved in reactions that divert resources away from the desired production pathway, or upregulating genes associated with the production pathway. Empirical designs based on detailed knowledge of the production and precursor pathways might not consider system-wide interactions and can, therefore, miss out on promising candidates in other parts of primary or secondary metabolism. Moreover, the nonlinear nature of metabolism makes it difficult to predict the systems-wide effects of simultaneous changes to multiple enzymes; this makes combinatorial design based on empirical knowledge challenging.

Computational models provide a systematic network-wide strategy for rational engineering by providing an efficient way to search the design space for suitable targets. Previous studies have relied on methods such as OptKnock^7^, optGene^8^, and optForce^9^ to successfully steer organisms like *S. cerevisiae* toward the overproduction of different compounds^10–14^. These methods use constraint-based models to predict what an overproducing phenotype should look like; however, they have limited capacity to predict actual cellular behavior because they are just a snapshot of the cellular state and lack kinetic information^15^. As a result, most of the targets suggested by these algorithms went through a vetting process to ensure they were not detrimental to the transient behavior of the cell. Moreover, these constraint-based approaches are ill-suited for developing combinatorial designs as they cannot capture the nonlinear epistatic interactions of simultaneous interventions^16^. Kinetic models overcome these shortcomings by capturing the dynamic response of the entire metabolic network to single or combinatorial genetic perturbations^17–21^. In this manner, they can tell us how to steer an organism from its reference phenotype towards overproduction. Despite these advantages, kinetic models have not been widely used for strain design. Indeed, constructing large-scale kinetic models is challenging due to their parametric and structural uncertainty^22^. Moreover, evaluating all possible combinations of genetic interventions that improve production is not trivial. To overcome these two hurdles, we recently developed NOMAD, a framework that employs nonlinear large-scale kinetic models for rational strain design^23^.

In this study, we used NOMAD to suggest strategies that drive *S. cerevisiae* to overproduce *p*-coumaric acid (*p*-CA), an essential ingredient in the food, pharmaceutical, and cosmetic industries^24–26^. *S. cerevisiae* is widely used as a host organism due to its ability to grow at large scale, robustness across various conditions, and availability of advanced genetic engineering tools^27,28^. We began with a *S. cerevisiae* strain (ST10284), already engineered for *p*-CA production through multiple gene knockouts and heterologous pathway additions^29^. Based on experimental physiological data, we built nine large-scale, multi-compartment kinetic models of ST10284 and used these models to identify combinatorial designs of three simultaneous enzyme targets that increase *p*-CA yield from glucose. We selected ten robust designs across all nine models and verified their performance in nonlinear batch fermentation simulations. Promoter swapping and plasmid-based overexpression strategies enabled experimental implementation of these designs. Eight out of the ten recombinant strains exhibited increased *p*-CA titers in batch fermentations, demonstrating the potential of kinetic models in guiding microbial engineering toward desired phenotypes.

## 2 Methods and Materials

### 2.1 Strains, media, and chemicals

The study utilized a previously developed *S. cerevisiae* strain ST10284 as the reference strain for building kinetic models and further engineering. This strain is a derivative of the CEN.PK113-7D strain^30^ that was engineered by incorporating heterologous pathways to produce *p*-CA from both L-tyrosine and L-phenylalanine. It includes numerous modifications designed to enhance flux within the pentose phosphate, shikimate, and aromatic amino acid biosynthetic pathways, as previously described^29^ (Supplementary Table S1 and Supplementary Figure S1).

Three kinds of medium were utilized in this study: YPD (yeast extract peptone dextrose), SC (synthetic complete dextrose), and MM (minimal medium). YPD medium is composed of 10 g/L yeast extract, 20 g/L peptone, and 20 g/L glucose. When required, we added 200 mg/L of G418 (G8168, Merck Life Science) and 100 mg/L of nourseothricin (Jena Bioscience, AB-101) as antibiotics for selection. SC medium is composed of 6.7 g/L yeast nitrogen base without amino acids, 1.4 g/L yeast synthetic drop-out medium supplement without histidine, leucine, tryptophan, and uracil, and 20 g/L glucose. The nucleotide uracil (76 mg/L, U1128, Sigma) was added when *URA3* was the selection maker for yeast transformation. MM is composed of per liter 7.5 g (NH_4_)_2_SO_4_, 14.4 g KH_2_PO4, 0.5 g MgSO_4_ 7H_2_O, 22 g dextrose, 2 mL trace metals solution (3.0 g/L FeSO_4_·7H_2_O, 4.5 g/L ZnSO_4_·7H_2_O, 4.5 g/L CaCl_2_·2H_2_O, 0.84 g/L MnCl_2_·2H_2_O, 0.3 g/L CoCl_2_·6H_2_O, 0.3 g/L CuSO_4_·5H_2_O, 0.4 g/L Na_2_MoO_4_·2H_2_O, 1.0 g/L H_3_BO_3_, 0.1 g/L KI, and 19.0 g/L Na_2_EDTA·2H_2_O), and 1 mL vitamins (0.05 g/L D-biotin, 0.2 g/L p-aminobenzoic acid, 1.0 g/L calcium D-pantothenate, 1.0 g/L thiamin–HCl, 1.0 g/L pyridoxine–HCl, 1.0 g/L nicotinic acid, and 25.0 g/L myo-inositol) with pH at 6. 2% bacteriological agar (A5306, Sigma) was added to the above media when preparing agar plates. All the standards used for analysis were purchased from Sigma.

The study used MM for the seed train and fed-batch cultivation of ST10284, as well as for the batch production cultivation of the engineered strains. For fed-batch cultivation, a 10-fold higher concentration of the components was utilized and the pH was adjusted to 5.6 using KOH to avoid the precipitation of salts. SC was used to assess the strength of the evaluated promoters, while YPD was used to select strains after yeast transformations. Glucose was the carbon source in all the cultivations.

### 2.2 Fed-batch cultivation conditions

The study utilized an established fed-batch cultivation setup to determine the parameters that characterized the phenotype of ST10284. To initiate the seed culture, ST10284 was streaked from −80 °C cryostocks onto YPD plates and incubated for 48 hours at 30 °C. A single colony was then inoculated into 1 mL MM containing 6 g/L of glucose and cultivated for 10 hours at 250 rpm. Cells were subsequently transferred to an initial OD_600_ of 0.1 across 8 wells of the fed-batch fermentation plate.

Fed-batch fermentation was performed in the microfluidic bioreactor Biolector Pro (m2p labs) using a 32 wells microfluidic flower plate (MTP-MF32C-BOH2), with each well containing 800 μL of batch MM supplemented with 1.29 g/L of glucose. We then filled two reservoir wells with 1800 μL of feed medium, with each reservoir feeding 4 cultivation wells, and triggered feeding after 18 hours using an exponential feeding profile, *F*(*t*) = 1.23 *e*^0.1t^, ensuring no overflow metabolism or oxygen limitation. Cultivation conditions were maintained at 30 °C, 1,300 rpm stirring speed, 35% oxygen (enriched air), and 85% humidity, with pH left uncontrolled during the whole cultivation. Glucose, biomass, *p*-CA, and ethanol concentrations were measured over approximately 7 hours, covering about one doubling period, with data collected from four samples at regular intervals.

### 2.3 Plasmid construction and strain construction

The chemical reaction targets suggested by NOMAD, along with all primers and biobricks used in this study, are summarized in Supplementary Tables S2–S4. *E. coli* competent cell *DH5α* was used for constructing and propagating all the plasmids corresponding to the different engineering strains (Supplementary Table S5). The *E. coli* strains grew at 37 °C on yeast extract tryptone (YT) medium with kanamycin (50 µg/mL), ampicillin (100 μg/mL), or chloramphenicol (25 ug/mL).

The strength of the promoters of the target genes was compared by amplifying the promoter region that was 1kb relative to the start codon location of the target genes using Phusion Hot Start II High-Fidelity PCR Master Mix (Thermo Scientific™, F565S) from *S. cerevisiae* genome. Afterwards, these amplified promoters, along with other selected promoters and the GFP coding sequence, were assembled into the pYTK096 vector^31^ using the Golden Gate assembly method as described in Shaw et al.’s work^32^. For gene knock-in, DNA fragments spanning 500 bp upstream and downstream of the target genes were amplified from the *S. cerevisiae* genome and fused with the corresponding weaker promoters via overlapping PCR to create repair templates. The guide RNA (gRNA) vectors targeting these genes were constructed using Gibson Assembly Master Mix (New England BioLabs).

The designs proposed by the kinetic models included both native targets from *S. cerevisiae* and heterologous targets introduced during the construction of ST10284. Phusion U Hot Start PCR Master Mix (Thermo Scientific™, F533S) was used to amplify the native genes from *S. cerevisiae*. For the heterologous targets, we amplified *EcAroB* and *EcAroD* from the plasmid pCFB1955^33^ and obtained *FjTAL* from Twist Bioscience (USA). The amplified target genes were integrated into ST10284 by cloning them into a set of vectors from the EasyClone 2.0 Yeast Toolkit using the USER cloning method^34^. All the constructed plasmids were subsequently verified by Sanger sequencing (Eurofins Scientific SE).

Recombinant strains (Table S6) were constructed using a Cas9-assisted approach^34^, followed by yeast transformations using the standard lithium acetate method^35^. Correct transformants were identified using the Phire Plant Direct PCR Master Mix (Thermo Fisher Scientific, F160S).

### 2.4 Fluorescence measurement

Single-cell fluorescence analysis was conducted using the NovoCyte Quanteon flow cytometry (Agilent). Yeast cultures (200 μL) were diluted by tenfold in a 96-well microtiter plate before analysis, with GFP fluorescence excited by a 488 nm laser and detected through the 525/50 nm channel. For each sample, we collected data from 10,000 cells and processed them using the FlowJo software.

### 2.5 Growth detection using Growth Profiler

Growth of the engineered yeast strains was monitored using the Growth Profiler 960 (EnzyScreen BV) at 30 °C with shaking at 250 rpm, according to the manufacturer’s recommended settings. Each experiment was conducted in triplicate, with images captured every 30 minutes over a 72-hour cultivation period using a 5ms shutter time and an initial OD600 of 0.2. G-values were extracted from the central pixel of each well using the GP960Viewer software and converted into OD_600_ based on established calibration curves.

### 2.6 Extraction and quantification of *p*-CA

Strains were cultivated for 5, 24, 30, 46, and 72 hours using the Growth Profiler 960. For *p*-CA analysis, 1 mL of yeast culture was mixed with 1mL of absolute ethanol, vortexed thoroughly for 10 seconds and centrifuged at 17,000xg for 10 minutes. An aliquot of the resulting supernatant was analyzed by high-performance liquid chromatography (HPLC) to determine the extracellular *p*-CA concentration. Additionally, 1 mL of yeast culture was centrifuged for 10 minutes, and the supernatant was used to quantify ethanol and glucose concentrations by HPLC.

For *p*-CA quantification, the HPLC system was equipped with a Discovery HS F5 column (150 mm × 2.1 mm, 3 μm particle size, Supelco, 567503-U). The column oven was set to 30 °C with a flow rate of 0.7 mL/min, and a 5 μL sample was injected. The mobile phase consisted of Solvent A (10 mM ammonium formate, pH 3.0, adjusted with formic acid) and Solvent B (acetonitrile). The initial solvent composition was 95% A and 5% B, maintained for 0.5 minutes, followed by a linear gradient to 40% A and 60% B at 7.0 minutes. This composition was held constant for 2.5 minutes (7.0–9.5 min) and then returned to the initial composition for the remainder of the run (9.6–12 min). *p*-CA was detected at a retention time of 4.7 minutes using absorbance at 277 nm. Glucose and ethanol were quantified using an Aminex HPX-87H column (Bio-Rad Laboratories, USA) under the conditions described by Borja et al.^36^, but with a column temperature of 60 °C.

### 2.7 Kinetic model construction

The NOMAD protocol was used to generate kinetic models representative of the *S. cerevisiae* strain ST10284^23^. We started with data integration, followed by the generation of steady-state profiles consistent with the integrated data using pyTFA^37^. Kinetic models were then generated around the steady-state closest to the mean of all the sampled states using the ORACLE^15,38^ framework implemented in the SKiMPy toolbox^39^. The generated parameter sets were then pruned for those that best represented the strain’s physiology.

#### 2.7.1 Model stoichiometry

We used redGEM^40^ to reduce the Yeast8 genome-scale model (GEM)^41^ around the central carbon metabolism subsystems: glycolysis, pentose phosphate pathway (PPP), cytosolic tricarboxylic acid cycle (TCA), mitochondrial TCA, shikimate pathway, and the anaplerotic reactions. Subsequently, the biosynthetic reactions were lumped into a single reaction depicting growth using lumpGEM^42^.

The model was then tailored to the strain ST10284 by incorporating its specific genetic modifications. Reactions corresponding to the heterologously expressed tyrosine ammonia lyase (*FjTAL*), phenylalanine ammonia lyase (*PAL2*), cinnamate-4-hydroxylase (*C4H*), fructose 6-phosphate kinase (*FPK*), and cytochrome P450 reductase 2 (*CYP5*) were added. The reaction PPYRDC corresponding to the knockout of *PDC5* was removed. The experimental strain ST10284 included the deletion of *Aro10*; however, we did not remove the two associated reactions in the model (INDPYRD and PYRDC) as other isoenzymes could catalyze them. We then included the reaction, FMN reductase (FMNRx), and transport reactions for acetyl phosphate and *p*-CA to ensure the system was mass-balanced. The resulting model had 368 reactions involving 295 metabolites distributed across nine compartments.

#### 2.7.2 Data integration

The pyTFA^37^ tool was used to integrate all metabolomic, fluxomic, and thermodynamic data as follows.

##### 2.7.2.1 Metabolomics

No data were available on the intracellular metabolite concentrations for strain ST10284; therefore, these concentrations were constrained to within a 5-fold range of the values reported in Park et al.^43^ for *S. cerevisiae*. For the extracellular metabolites, we used data from the ST10284 fermentation experiments. The concentrations of oxygen, glucose, and other components of the mineral medium were constrained to be within 20% of their experimental values. The extracellular pH was set at 6, the starting pH value in fermentation experiments. The remaining extracellular metabolites that were not detected were constrained to a maximum concentration of 1 μ*mol*. Finally, the concentrations of the components of the cytosolic adenylate pool were constrained to ensure that the adenylate energy charge was sufficient^44^.

##### 2.7.2.2 Fluxomics

Exofluxomics data from ST1084 fermentations were used to constrain the growth, secretion, and uptake rates in the model. The p-CA secretion rate was constrained within the experimentally observed range around the mean, while ethanol secretion was constrained between the experimentally observed mean and one standard deviation below the mean to enforce selectivity toward p-CA production. The glucose uptake rate was allowed to vary within 20% and 10% of the upper and lower experimental bounds, respectively. Growth was constrained to be at least the lower bound of the experimentally reported range (Supplementary Table S6). Additionally, the ratio of the fluxes through glucose-6-phosphate isomerase (PGI) and glucose 6-phosphate dehydrogenase (G6pDH2r) was constrained to be consistent with the physiology of *S. cerevisiae*^45,46^.

##### 2.7.2.3 Thermodynamics

Thermodynamics-based flux balance analysis (TFA) uses thermodynamics data to ensure that the sampled fluxes and concentrations adhere to the second law of thermodynamics. To this end, we used data from the equilibrator API^47^ to impose bounds on the standard Gibbs free energy (Δ*G*^’0^) for the different reactions in the model. Reactions without thermodynamic data were constrained to have their Δ*G*^’0^ = 0 ± 2 *kcal*/*mol*. Furthermore, intercompartmental transport reactions were constrained to operate close to thermodynamic equilibrium.

Some reactions could carry fluxes in both directions even after adding the different omics data. We manually curated the directions for these reactions; for example, we forced the TCA to be cyclic and the pentose phosphate pathway to have a catabolic non-oxidative branch^48^.

#### 2.7.3 Kinetic models consistent with integrated data

We assigned a kinetic mechanism to each reaction in the model (Supplementary Data 1) and generated mass balance equations for all metabolites, excluding inorganic salts. The resulting system of ordinary differential equations (ODEs) consists of 268 mass balances, including one for biomass, across 303 reactions, parameterized by 1098 Michaelis constants (*K*_M_) and 303 maximal reaction velocities (*V*_max_*s*). Volume fractions were assigned for all the compartments; four out of the nine contained at least one mass balance that was not a proton - the cytosol (70%), mitochondria (3%), endoplasmic reticulum (2%), and the endoplasmic reticulum membrane (0.5%) (Supplementary Data 1).

With the model structure in place, the next task was to parameterize it. Experimental data on kinetic parameters are limited, and reported values can vary substantially. For example, BRENDA^49^ entries were available for only 164 out of the 1098 *K*_M_ in our model, and 15 of these *K*_M_ had values spanning more than two orders of magnitude. Moreover, the set of kinetic parameters consistent with the observed steady state can be non-unique, i.e., multiple sets of parameters can align to the same steady-state conditions. NOMAD accounts for this parametric uncertainty by identifying a *population* of kinetic models that are most likely to represent the strains’ true physiology rather than relying on a single model.

Following the ORACLE framework, we initially generated a putative set of models as follows. Using pyTFA, we sampled 5,000 steady states that produced at least 95% of the maximal growth under the applied constraints. The kinetic models were constructed around the steady-state profile that was closest to the mean of these samples. We constrained *K*_M_values to fall within the minimum and maximum values reported in BRENDA^49^ for *S. cerevisiae* across all reactions. We then sampled 15,000 sets of kinetic parameters that were consistent with the steady-state profile and resulted in locally stable models.

In addition to local stability checks^50^, NOMAD uses a two-step process to identify models that capture not only the steady-state behavior of the strain but also its dynamic properties. In the first step, models were screened for those with dominant time constants at least three times smaller than the doubling time of the strain (7.3 hours). This criterion is based on the timescales involved in metabolism and the likelihood of the organism returning to its homeostatic state after perturbation^18,20,23^. The models that passed this step were further pruned by selecting those that exhibited robust growth in batch fermentation simulations. The resulting models are likely to accurately represent both the steady-state and dynamic characteristics of the strain under fermentation conditions. We quantified the variation across these models by calculating the coefficient of variation (CV), defined as the ratio of the standard deviation to the mean, for each of the 1098 *K*_#_*s* and 303 *V*_max_*s*.

The NOMAD protocol outlined above used steady-state data from fed-batch fermentation experiments to build the models. However, the final design strains would be implemented in a batch setting, where yeast exhibits a 3-fold increase in growth rate compared to the fed-batch conditions. We accounted for this altered phenotype by applying a 3-fold increase to the maximal velocities of all the reactions in the chosen models while keeping the parameters such as the Michaelis constants, thermodynamic constants, and the initial conditions fixed. We validated this approach by conducting a sensitivity analysis on kinetic models built around steady states constrained to have a 3-fold increase in exofluxomics and a 20% variability in the internal concentrations. This analysis showed that the distribution of maximal velocities in the resulting kinetic models had a median 3-fold increase compared to the models calibrated on fed-batch data, suggesting that this approach accurately captures the altered metabolic phenotype (Supplementary Note I and Supplementary Figure S2).

### 2.8 Rational design generation for overproducing *p*-CA

Kinetic models enable the calculation of control coefficients, which quantify how a change in an enzyme activity or metabolite concentration impacts fluxes and concentrations throughout the network^51,52^. Network Response Analysis (NRA) makes use of these control coefficients to efficiently generate combinatorial designs based on log-linear approximations of the kinetic models^53^. We used NRA to identify engineering strategies to maximize the yield of *p*-CA from glucose while considering network-wide effects. Strategies adhered to the following constraints: (i) maximum 2-fold changes in enzyme activity, (ii) no more than 5-fold changes in concentrations, (iii) no more than 3 simultaneous target enzymes, and (iv) maximum 20% reduction in growth rate. Considering the extensive engineering of the reference strain ST10284, the design constraints were informed by the feasibility of experimental implementation and validation. Specifically, the number of target enzymes was limited to three to balance the exploration of metabolic design space with the practical constraints of strain construction and phenotypic characterization. Introducing more targets could increase experimental complexity and the risk of unintended metabolic interactions. The constraint on permissible fold changes in enzyme activity was based on prior experience with reliably achievable over- and under-expression levels. Similarly, the concentration fold-change constraint was determined based on prior modeling experience^23^, ensuring that the proposed perturbations remained within biologically realistic and experimentally achievable ranges. Together, these constraints enabled a focused yet feasible strain optimization strategy.

For each kinetic model, we first maximized the yield of *p*-CA while adhering to the design constraints. We then generated combinations of 3 enzymes that produced *p*-CA yield increases were within 5% of the maximal predicted yield.

#### 2.8.1 Robustness evaluation

The population of models can span a wide range of parameter values. Therefore, designs generated with one model need not perform well across other models. We accounted for this parametric uncertainty by identifying designs that produce robust performance across the population of models. NRA was chosen for this task due to its computational efficiency, which arises from the use of log-linear approximations of kinetic models based on their control coefficients^51,52^. For each combination of design and model, we maximized the yield of *p*-CA from glucose while permitting changes to only the 3 enzymes in the design. This procedure resulted in a matrix of predicted yield increases for each combination of model and design. We then ranked the designs based on the mean of these predicted yield increases and chose the top 10 for further analysis and experimental implementation.

#### 2.8.2 Verification through nonlinear simulations

We verified the log-linear predictions by implementing the top 10 NRA-predicted designs in full-scale nonlinear simulations and calculating the relative increase in *p*-CA yield from glucose when compared to the reference strain, ST10284.

For each combination of model and design from the Top 10, we adjusted the maximal velocities of the associated enzymes according to the NRA-suggested values. Batch fermentation was then simulated for 48 hours, starting with an inoculum of 0.024 g/L and using the same medium conditions applied during kinetic model development. The yield of *p*-CA from glucose and the *p*-CA titer were recorded at 5 hours and 24 hours, respectively, and their increase relative to ST10284 was calculated.

## 3 Results

### 3.1 Kinetic models of a *p*-CA producing *S. cerevisiae* strain

We followed the NOMAD protocol established by Narayanan et al^23^ to build a population of kinetic models representing *S. cerevisiae* ST10284 physiology (Supplementary Table S6) that can effectively simulate the strain’s behavior in batch reactor conditions. The framework identified nine models that produced adequate growth in batch fermentation. The models captured the behavior of 257 metabolites and biomass connected by 303 reactions across the central carbon pathways such as glycolysis, the pentose phosphate pathway, the Krebs cycle, the Shikimate pathway, and the electron transport chain. The nine models covered a wide range of parameter values that could manifest in the observed phenotype of the reference strain, with the coefficient of variation (Methods) ranging from 41% to 295% and 31% to 299% for the 1098 *K*_M_ and 303 *V*_max_*s*, respectively (Supplementary Data 1). Of the 164 *K*_M_values available in BRENDA for *S. cerevisiae*, 93 overlapped with those from the nine models, while the nine models’ values for the remaining 36 fell within an order of magnitude of the experimental ranges. The model parameters demonstrated robustness in nonlinear settings, successfully simulating growth and p-CA production in batch fermentation.

### 3.2 Model-based designs for overproducing *p*-CA in *S. cerevisiae*

We applied NRA to the nine kinetic models to devise designs for improved yield of *p*-CA from glucose in the *p*-CA producing strain, ST10284. Each design strategy was limited to three target enzymes, with up to a 2-fold increase in their activity and unlimited downregulation (Methods). To minimize disruption to other cellular functions, metabolite concentration changes were restricted to a 5-fold difference. Designs were selected based on their ability to achieve a p-CA yield increase within 5% of the model’s maximal yield.

Across the nine kinetic models, we identified 39 unique designs, each defined by a distinct set of three target enzymes. Clustering analysis grouped these designs into four broad clusters based on the similarity of their enzyme targets (Figure 1). Group I contains two subgroups of two designs each; one subgroup targets glutamate dehydrogenase (GLUDy) and isocitrate dehydrogenase (ICDHyr) together, while the other targets sedoheptulose 1,7-bisphosphate D-glyceraldehyde-3-phosphate-lyase (FBA3). The 17 designs in Group II all increase the availability of phosphoenolpyruvate (pep), a precursor to the aromatic amino acid pathways, by decreasing its consumption by pyruvate kinase (*PYK*). Substitution of the PYK promoter has previously been shown to increase *p*-CA production in *S. cerevisiae*.^54^ Ten out of the 17 designs also increase activity through the production pathway by upregulating either the heterologous tyrosine ammonia lyase (TAL in our model), prephenate dehydrogenase (PPND2), or 3-dehydroquinate dehydratase (DHQTi). Groups III and IV focus on the *p*-CA production and shikimate pathways, respectively. All seven designs in group III suggest upregulating TAL, PPND2, or aspartate transaminase (ASPTA), with four designs containing an additional target among the three. In group IV, all nine designs target 3-dehydroquinate synthase (DHQS), with seven also upregulating chorismate mutase (CHORM).

**Figure 1:**
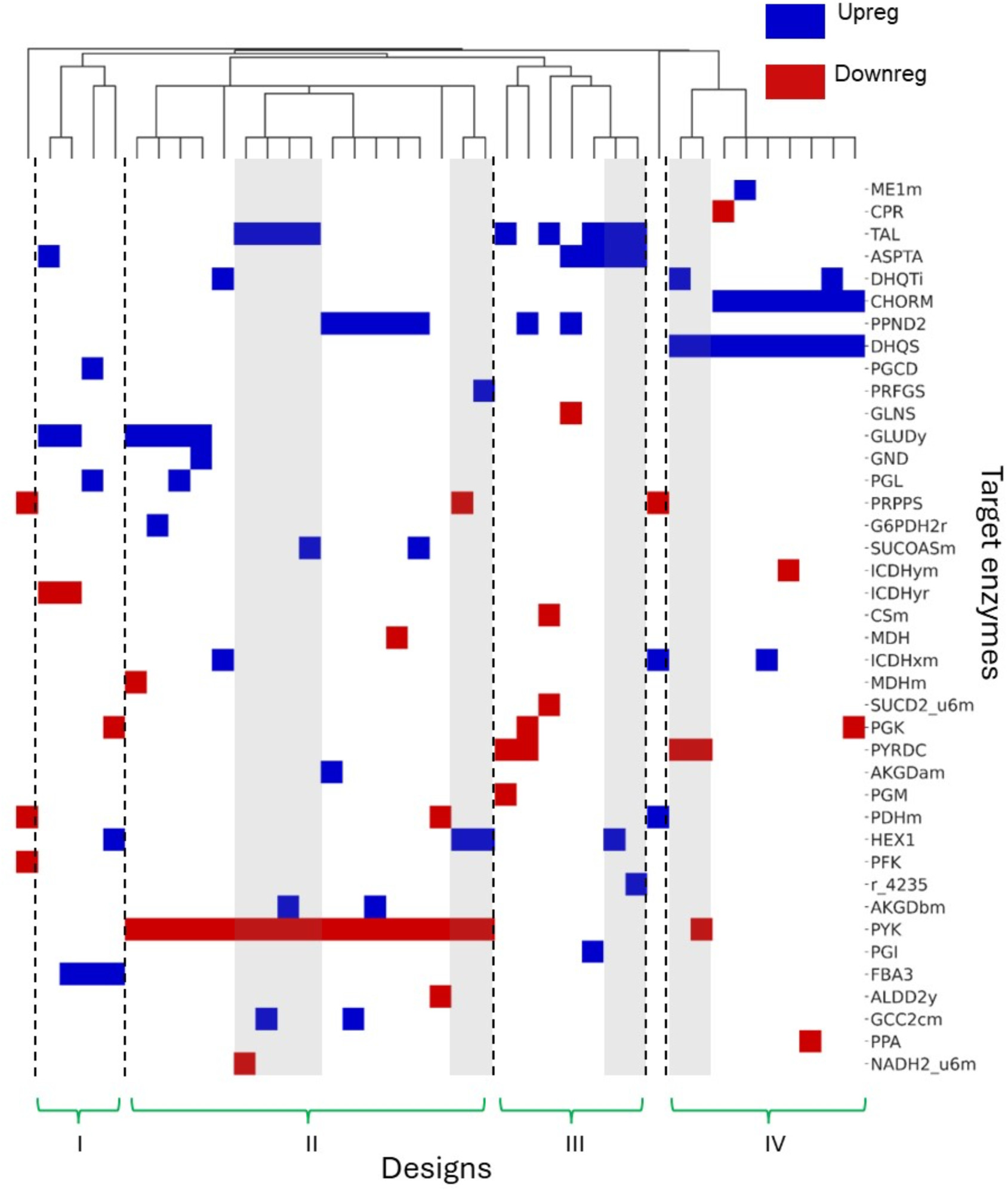
Clustering analysis of the 39 unique designs (columns) generated using NRA and their associated enzymes (rows). Each column represents the three upregulated (blue) or downregulated (red) enzymes targeted by a single design. The target enzymes are contained mainly in the pentose phosphate pathway (PPP), the Krebs cycle, glycolysis, and the p-CA production pathway. Six of the 10 experimentally implemented designs (gray-shaded columns) lie within cluster II. The remaining 4 designs belong to clusters III and IV. **AKGDam:** Oxoglutarate dehydrogenase lipoamide, **AKGDbm:** Oxoglutarate dehydrogenase dihydrolipoamide S succinyltransferase, **ALDD2y:** Aldehyde dehydrogenase, **ASPTA:** Aspartate transaminase, **CHORM:** Chorismate mutase, **CPR:** Cytochrome P450 reductase 2, **CSm**: Citrate synthase, **DHQS:** 3-dehydroquinate synthase, **DHQTi:** 3-dehydroquinate dehydratase irreversible, **FBA3:** Sedoheptulose 1,7-bisphosphate D-glyceraldehyde-3-phosphate-lyase, **GCC2cm:** Glycine cleavage complex lipoamide mitochondrial, **GLNS:** Glutamine synthetase, **GLUDy:** Glutamate dehydrogenase, **GND:** Phosphogluconate dehydrogenase, **G6PDH2r:** Glucose 6-phosphate dehydrogenase, **HEX1**: Hexokinase, **ICDHxm:** Isocitrate dehydrogenase, **ICDHym:** Isocitrate dehydrogenase mitochondrial, **ICDHyr:** Isocitrate dehydrogenase, **MDH:** Malate dehydrogenase, **MDHm:** Malate dehydrogenase mitochondrial, **ME1m:** Malic enzyme mitochondrial, **NADH2_u6m:** NADH dehydrogenase mitochondrial, **PDHm:** Pyruvate dehydrogenase, **PFK:** Phosphofructokinase, **PGCD:** Phosphoglycerate dehydrogenase, **PGI:** Glucose-6-phosphate isomerase, **PGK:** Phosphoglycerate kinase, **PGL:** 6-phosphogluconolactonase, **PGM:** Phosphoglycerate mutase, **PPA:** Inorganic diphosphatase, **PPND2:** Prephenate dehydrogenase, **PRFGS:** Phosphoribosylformylglycinamidine synthase, **PRPPS**: Phosphoribosylpyrophosphate synthetase, **PYK:** Pyruvate kinase, **PYRDC:** Pyruvate decarboxylase, **r_4235:** Glucokinase paralog EMI2, **SUCD2_u6m:** Succinate dehydrogenase ubiquinone 6 mitochondrial, **SUCOASm:** Succinate CoA ligase, **TAL:** Tyrosine Ammonia Lyase (heterologous).

We ranked these 39 designs based on their mean NRA predicted performance across the models (Methods). The top 10 designs (Figure 1, gray shaded columns; Table 1) all predicted a mean increase of 12-14% in the *p*-CA yield from glucose relative to ST10284. Cluster II accounted for six of these designs, which were further categorized into two subgroups.

**Table 1:** Top 10 NOMAD designs implemented experimentally. Sets of 3 enzymes targeted by each design, along with their reaction names in the model, the genes targeted experimentally, and the name of the corresponding recombinant strain. We implemented design 7 without SUCOASm as implementing it experimentally caused yeast death.

| Design | Reaction | Enzyme | Genes | Strain |
| --- | --- | --- | --- | --- |
| 1 | HEX1↑<br>PRPPS↓<br>PYK↓ | Hexokinase<br>Phosphoribosylpyrophosphate synthetase<br>Pyruvate kinase | *HXK2*<br>*PRS1*<br>*PYK1* | ST14204 |
| 2 | HEX1↑<br>PYK↓<br>PRFGS ↑ | Hexokinase<br>Pyruvate kinase<br>Phosphoribosylformylglycinamide synthase | *HXK2*<br>*PYK1*<br>*ADE6* | ST14210 |
| 3 | PYK↓<br>AKGDbm↑<br>TAL↑ | Pyruvate kinase<br>Mitochondrial 2-oxoglutarate dehydrogenase complex<br>Tyrosine ammonia-lyase | *PYK1*<br>*KGD1*, *KGD2*, *LPD1*,<br>*YMR31*<br>*FjTAL* | ST14206 |
| 4 | PYK↓<br>TAL↑<br>GCC2cm↑ | Pyruvate kinase<br>Tyrosine ammonia-lyase<br>Glycine decarboxylase multienzyme complex | *PYK1*<br>*FjTAL*<br>*GCV1*, *GCV2*, *GCV3*,<br>*LPD1* | ST14207 |
| 5 | HEX1↑<br>TAL↑<br>ASPTA↑ | Hexokinase<br>Tyrosine ammonia-lyase<br>Aspartate transaminase | *HXK2*<br>*FjTAL*<br>*AAT2* | ST14186 |
| 6 | PYK ↓<br>DHQS↑<br>PYRDC↓ | Pyruvate kinase<br>3-dehydroquinate synthase<br>Pyruvate decarboxylase | *PYK1*<br>*EcAroB*<br>*PDC1* | ST14211 |
| 7 | PYK↓<br>TAL↑<br>SUCOASm↑ | Pyruvate kinase<br>Tyrosine ammonia-lyase<br>Mitochondrial succinyl-CoA synthetase complex | *PYK1*<br>*FjTAL*<br>*LSC1*, *LSC2* | ST14201 |
| 8 | DHQTi↑<br>DHQS↑<br>PYRDC↓ | 3-dehydroquinate dehydratase<br>3-dehydroquinate synthase<br>Pyruvate decarboxylase | *EcAroB*, *EcAroD*<br>*EcAroB*<br>*PDC1* | ST14212 |
| 9 | PYK↓<br>TAL↑<br>NADH2_u6m↓ | Pyruvate kinase<br>Tyrosine ammonia-lyase<br>NADH dehydrogenase mitochondrial | *PYK1*<br>*FjTAL*<br>*NDE1* | ST14195 |
| 10 | TAL↑<br>ASPTA↑<br>r_4235↑ | Tyrosine ammonia-lyase<br>Aspartate transaminase<br>Glucokinase | *FjTAL*<br>*AAT2*<br>*EMI2* | ST14205 |

The first subgroup, consisting of four designs, suggested increasing flux through the production pathway by upregulating TAL while simultaneously increasing the availability of pep for the shikimate pathway through PYK downregulation. Three of the four designs included a third target in the TCA cycle (GCC2cm/AKGDbm/SUCOASm) that could ensure sufficient energy for growth. The remaining design contained a counterintuitive target - the downregulation of NADH dehydrogenase (NADH2_u6m), which serves as the point of entry to the electron transport chain.

The second subgroup ensured sufficient precursor availability for *p*-CA production by simultaneously upregulating hexokinase (HEX1) and downregulating PYK. The third suggested intervention was either the upregulation of phosphoribosylformylglycinamidine synthase (PRFGS) or the downregulation of phosphoribosylpyrophosphate synthetase (PRPPS). The former could increase the availability of glutamate for tyrosine synthesis and subsequent *p*-CA production, while the latter could deprive anthranilate phosphoribosyltransferase (ANPRT) of its substrate prpp (5-Phospho-alpha-D-ribose 1-diphosphate), thereby reducing the redirection of carbon away from tyrosine/phenylalanine synthesis.

The remaining four designs were equally split between clusters III and IV. The designs in cluster III focused on increasing flux through the production pathway (ASPTA + TAL), while ensuring sufficient uptake of glucose through HEX1 or glucokinase encoded by the gene *EMI2* (r_4235 in our model). In cluster IV, the designs aimed to increase flux through DHQS in the shikimate pathway, while reducing pyruvate consumption by pyruvate decarboxylase (PYRDC). This is pertinent because increased pyruvate consumption would necessitate the redirection of pep towards pyruvate production. The third target in this cluster either aimed at complementing one of the other two modifications by further increasing the flux through the production pathway (DHQTi) or increasing the availability of pep through PYK downregulation.

We simulated the top 10 designs in a batch fermentation setting to evaluate their performance in a nonlinear context. Most design-model combinations increased the *p*-CA yield, with a maximum increase of 210%. Eight out of the top 10 designs demonstrated improvement in all nine models, while the remaining two designs did so for eight out of the nine models (Figure 2A). This yield improvement aligns with the NRA predictions, strengthening confidence in the experimental viability of these designs. Moreover, 39 out of the possible 90 combinations of design and model provided an increase in overall *p*-CA titer after 24 hours in batch simulations (Figure 2B), despite this not being an explicit design objective. For these 39 combinations, the engineered strains exhibited glucose uptake dynamics and overall consumption similar to the reference strain (Supplementary Figure S3); this observation was consistent with the design objective of improving *p*-CA yield. The increased *p*-CA production was accompanied by a slight reduction in mean biomass titers relative to ST10284 (Figure 2C).

**Figure 2:**
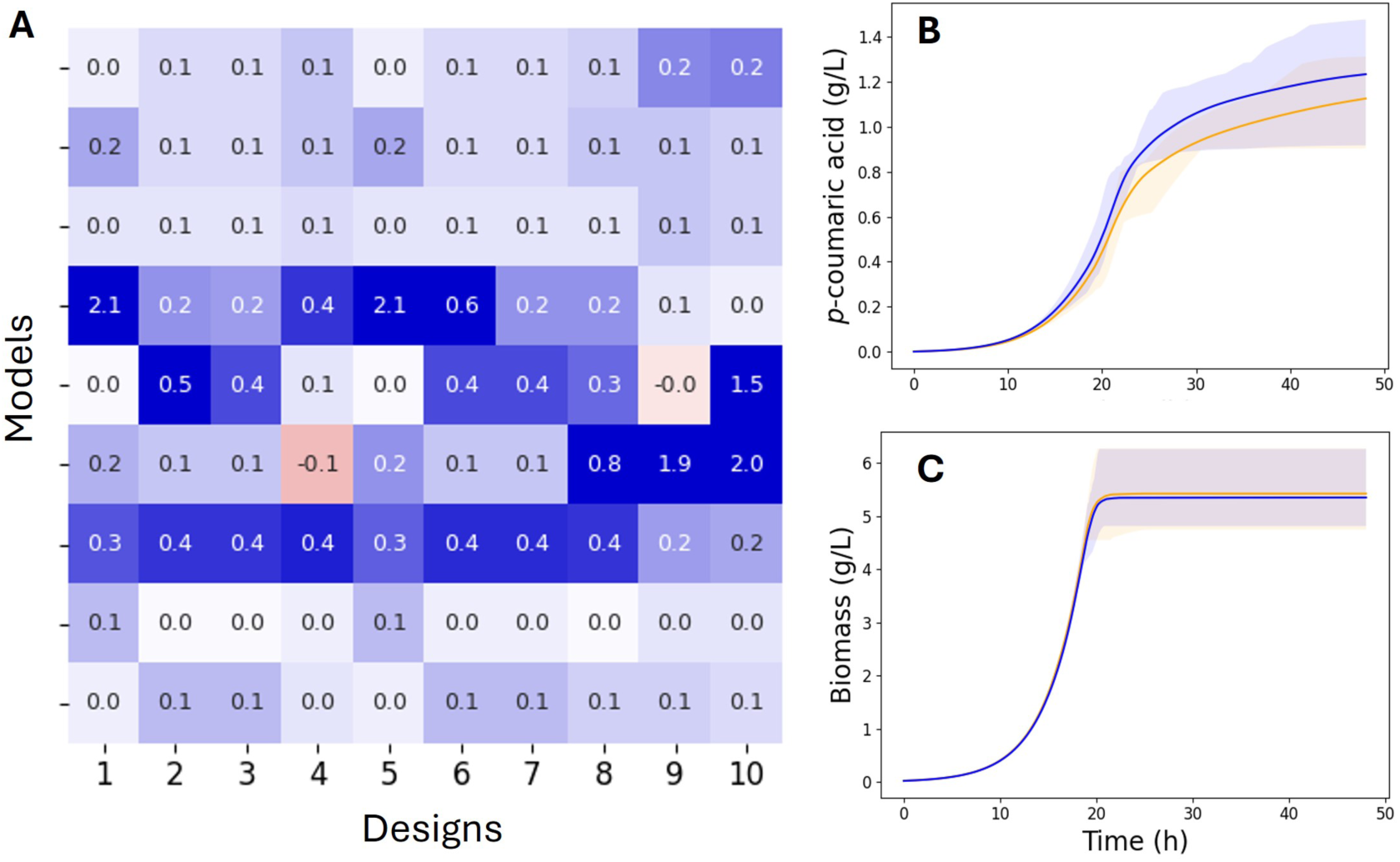
Performance of the top 10 designs evaluated in nonlinear batch simulations. (A) Relative increase (blue) and decrease (red) in the p-CA yield from glucose for each combination of design and model, compared to ST10284. The yield was calculated in the mid-exponential phase, at t=5 hours. All 10 designs produced a higher yield for 8 out of 9 models, while 8 designs produced a higher yield for all 9 models. (B, C) Simulated p-CA and growth titers for the reference strain (orange) and the top 10 designs (blue) across the 39 combinations of models and designs that provided an increase in p-CA titer after 24 hours. The solid line is the mean across the models, and the shaded region represents the interquartile range.

### 3.3 Reconstruction of rationally engineered *S. cerevisiae* strains

The 10 designs (Table 1) suggested by the kinetic models required several simultaneous up and down regulations of enzyme activities. We first implemented strategies to downregulate the activities of pyruvate kinase (PYK, designs 1-4, 6, 7, and 9), phosphoribosylpyrophosphate synthetase complex (PRPPS, design 1), NADH dehydrogenase (NADH2_u6m, design 9), and pyruvate decarboxylase (PYRDC, designs 6 and 8). These enzymes all play pivotal roles in the homeostatic function of the strain. Pyruvate kinase has paralogs *PYK1* and *PYK2 and* has been reported as the primary enzyme in the glycolytic pathway^55,56^. NADH dehydrogenases *NDE1* and *NDE2* play roles in transferring cytosolic NADH to the mitochondrial respiratory chain^57,58^. PRPPS is crucial for the synthesis of nucleotides, with the *PRS1* subunit being key in enzyme activity encoding in *S. cerevisiae*^59^. PYRDC is encoded by three isozymes - *PDC1*, *PDC5*, and *PDC6*, with PDC1 being the major isozyme^60^. We used a promoter-swapping strategy to implement these four downregulations using the key genes for each enzyme - PYK1, NDE1, PRS1, and PDC1. We first evaluated the strength of various promoters compared to the native promoters of these target genes, considering the −1kb region upstream of the start codon as the native promoter region. We linked all the promoters with GFP using Golden Gate Assembly and measured their expression levels by flow cytometry. The native promoters *PrNDE1* and *PrPRS1* were already relatively weak, with only *PrRAD27* and *PrREV1* providing a lower fluorescence intensity (Figure 3). The weaker of the two, *PrREV1,* was chosen as their replacement. Conversely, the native promoters *PrPYK* and *PrPDC1* were stronger than all the candidates (Figure 3C). Therefore, they were replaced with their weakest compatible counterparts - *PrPRL18b* and *PrTEF*, respectively.

**Figure 3:**
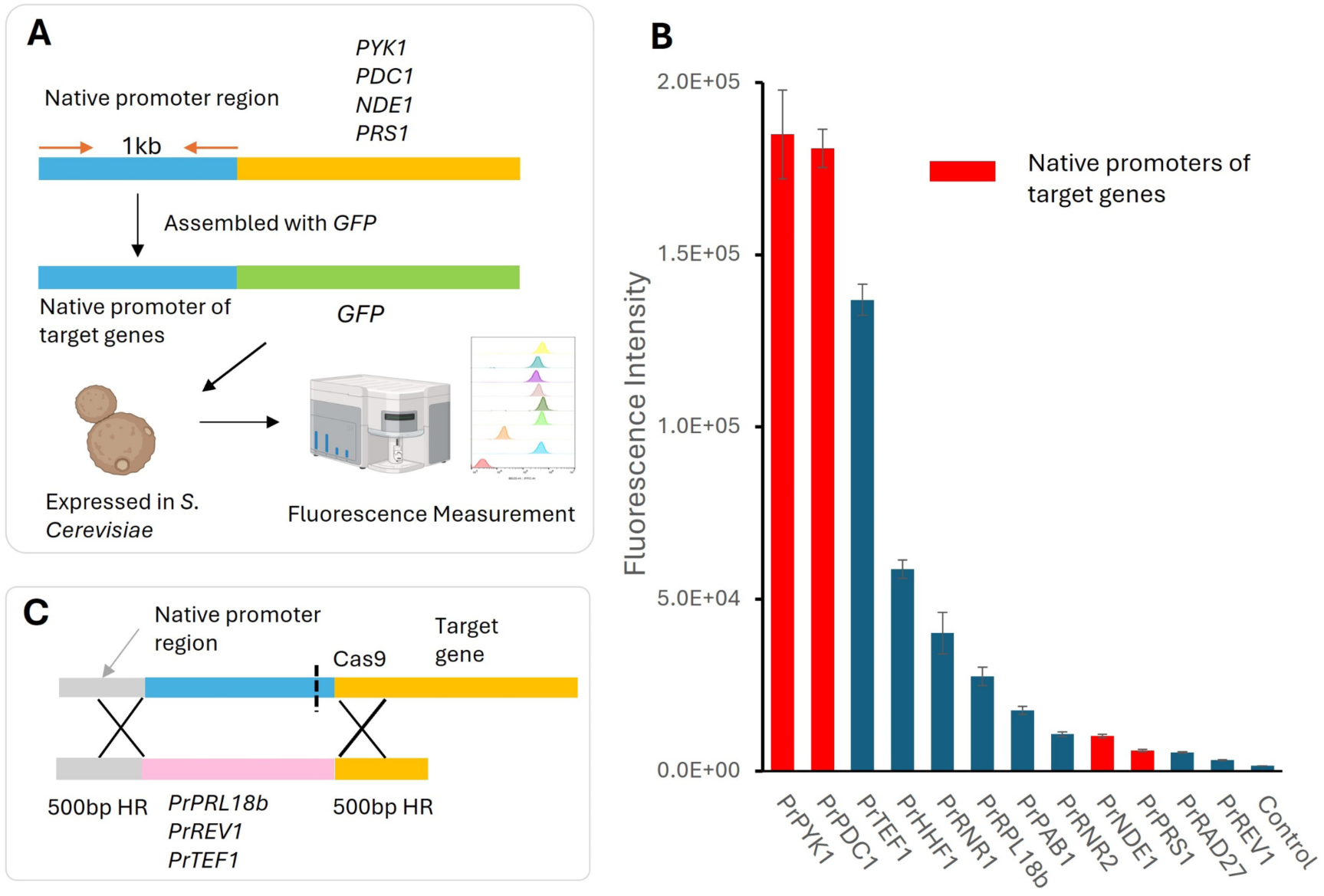
Gene downregulation through a promoter swapping strategy. **(A)** Design for measuring the strength of the promoter of four target genes PYK1, PDC1, NED1 and PRS1. **(B)** Promoter strength measured by flow cytometry. **(C)** Downregulation of the four target genes through swapping of the native promoter with a weaker promoter using CRISPR-aid.

Next, we implemented the strategy for upregulating the design targets suggested by the kinetic models. Designs 3, 5, 7, 9, and 10 involved the upregulation of tyrosine ammonia-lyase, corresponding to TAL from *Flavobacterium johnsoniae* (FjTAL). Three of the targets were enzyme complexes formed by multiple subunits –the glycine decarboxylase multienzyme complex (GCC2cm, design 4), the mitochondrial succinyl-CoA synthetase complex (SUCOASm, design 7), and the mitochondrial 2-oxoglutarate dehydrogenase complex (AKGDbm, design 3). We chose to upregulate all the subunits involved in these complexes as upregulating only some of the subunits might not produce a significant effect. We upregulated 3-dehydroquinate synthase (DHQS, designs 6 and 8) and 3-dehydroquinate dehydratase (DHQTi, design 8) using the *E. coli* genes *EcAroB* and *EcAroD* instead of the native pentafunctional gene *ARO1* as this enabled us to target them specifically. For hexokinase (*HEX1*, designs 1, 2, and 5) and aspartate transaminase (ASPTA, designs 5 and 10), we used their dominant paralogs, *HEX2* and *AAT2*, as targets for upregulation. *HEX2* is the predominant isoenzyme for hexokinase during the growth on glucose, playing the central role in glucose phosphorylation^61^. Similarly, *AAT2* encodes the cytosolic form of aspartate transaminase, the major contributor to total aspartate aminotransferase activity^62^. Finally, we used *S. cerevisiae*’s native gene *ADE6* for upregulating the activity of phosphoribosylformylglycinamidine synthase (PRFGS, design 2) and the gene *EMI2* for glucokinase (r_4235, design 10). We cloned all the target genes specific to each design into EasyClone vectors using USER cloning and expressed these targets in ST10284 using the CRISPR editing method. All designs could be implemented exactly except for design 7 (ST14201); we had to remove the overexpression of mitochondrial succinyl-CoA synthetase (SUCOASm) as it caused yeast death.

### 3.4 *p*-CA production in reconstructed strains

We tested the production capabilities of the recombinant strains (Table 1) in microplate batch fermentations using the Growth profiler shaker incubator. During the first 24 hours, all 11 strains – the reference strain and the 10 design strains – grew on glucose and produced ethanol (Supplementary Figures S4 and S5); these are the conditions under which we built the kinetic models and generated the *in-silico* designs. The strains then consumed ethanol for further growth and *p*-CA production. All the engineering strains, except ST14207 and ST14204, increased *p*-CA titers by 19-32% compared to the reference strain at the end of fermentation (Figure 4). ST14206 demonstrated the highest increase in titer (32%), while ST14210 provided the smallest increase (19%). These eight strains also produced more *p*-CA than the reference strain while growing on glucose alone, with increases ranging from 0.4% to 20% at 24 hours. Notably, all ten strains maintained a growth profile comparable to that of ST10284 during this period of growth on glucose; they attained biomass titers that were at least 85% of the reference strain at 24 hours. This adherence to the phenotypic behavior of the reference strain is pertinent as we explicitly constrained the generated designs to have minimal perturbation from the reference strain while satisfying the design objective. The eight strains that increased *p*-CA production maintained biomass production and reached final OD_600_ values of at least 90% of the reference OD_600_ at 72 hours. The two strains that did not exhibit increased *p*-CA titers, ST14204 (design 1) and ST14207 (design 4), reached substantially lower OD_600_ values. ST14207 maintained its growth and *p*-CA production during growth on glucose; however, it could not consume ethanol to fuel further growth and production (Supplementary Figure S4). ST14204, on the other hand, demonstrated impaired performance throughout the fermentation process. This is possibly due to the downregulation of PRPPS, which disrupts the synthesis of nucleotides, which are the fundamental building blocks of the cell.

**Figure 4:**
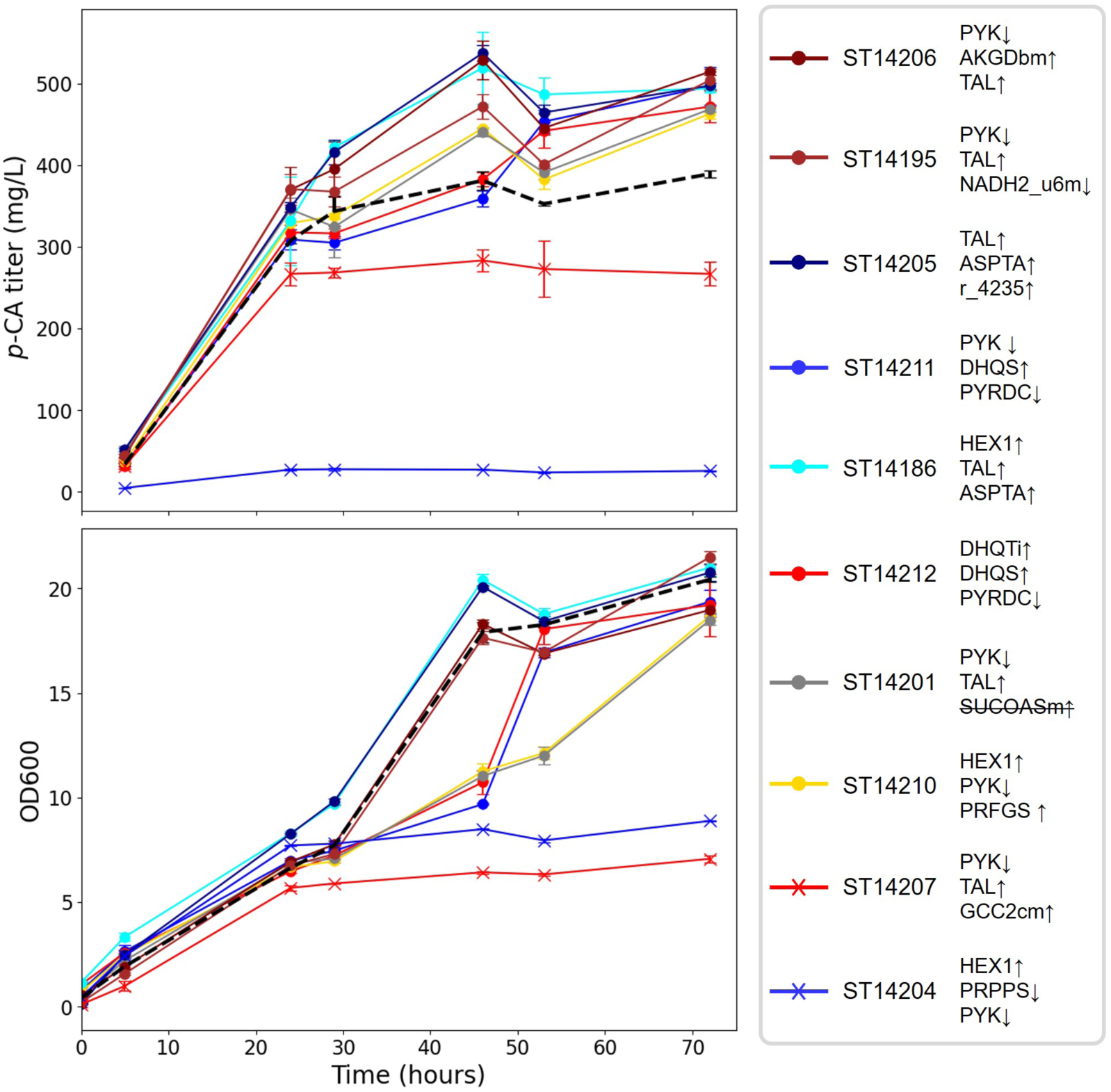
Average p-CA titers (top) and OD600 (bottom) of all ten engineered strains and the reference strain (dashed black) in triplicate batch fermentation experiments. The strains in the legend are in descending order of the recorded p-CA titer at 72 hours. All ten strains adhered to the growth dynamics when growing on glucose; they produced at least 85% of the biomass concentration of ST10284 at 24 hours. Eight out of the ten strains (solid dots) produced higher p-CA titers than ST10284 during both growth phases. They displayed p-CA titer increases of 0.4-20% and 19-32% when compared to the reference strain at 24 hours and 72 hours respectively. The two strains that did not improve p-CA titers were ST14207 (red cross) and ST14204 (blue cross). The error bars for all the strains represent the standard deviation across 3 repeats. The individual growth and p-CA production curves for the strains are provided in Supplementary Figure S6.

## 4 Discussion

Advances in genetic engineering techniques have enabled multiple enzymatic interventions towards improved cellular performance. This work showed how model-guided strain design can successfully be integrated into a design-build-test-learn cycle. We used the NOMAD framework to devise strategies for improved *p*-CA production in *S. cerevisiae*. Our kinetic-model-based combinatorial designs achieved a high success rate, with eight out of ten designs (80%) producing at least an 18% increase in *p*-CA titers at the end of fermentation.

This successful translation of in-silico designs into viable and high-performing strains was due to the use of (i) large-scale kinetic models that accurately capture the dynamic behavior of the reference strain, (ii) a constraint-based design that maintains the engineered strain’s phenotype close to that of the reference strain, and (iii) experimental strategies capable of fine-tuning gene expression levels.

Large-scale kinetic models capture the network-wide effects of genetic interventions, including their impact on key cellular health indicators, such as the energy charge and redox potential. We selected the models most likely representing the reference strain by using nonlinear simulations and concepts from control theory^23^. With these models, we used NOMAD to identify the top 10 designs among 4.5 million combinations of three simultaneous enzyme modifications that could reliably increase *p*-CA production, allowing us to save time and resources. These models also enabled us to identify non-obvious, off-pathway targets, such as NADH dehydrogenase and 2-oxoglutarate dehydrogenase in the mitochondrion, out of 303 possible genetic interventions and their combinations.

NOMAD further ensured the viability of proposed designs by limiting how much the designed strains could deviate from the reference strain. This phenotypic constraint proved essential; we observed that the successful designs also maintained at least 90% of the growth of the reference strain at the end of fermentation. Moreover, we achieved an 80% success rate by choosing only those designs that remained robust across the phenotypic variation covered by the population of nine kinetic models. The success of our top 10 designs also relied on their precise experimental implementation as they included targets such as PYK and NADH2_u6m that are critical to the homeostatic functioning of the strain. To tackle this challenge, we employed a promoter-swapping strategy for downregulations and overexpressions for upregulation.

While most of the designs translated well into improved strains, two—ST14204 and ST14207—did not yield the desired outcomes. ST14204, which involved upregulation of HEX1 and downregulation of PYK and PRPPS, exhibited relatively good growth but almost no p-CA production. The repression of PRPPS (phosphoribosyl pyrophosphate synthetase), a key enzyme in generating PRPP and feeding the shikimate pathway, likely restricted precursor supply for aromatic amino acid and *p*-CA biosynthesis. This hypothesis is supported by the success of ST14210, which shared targets HEX1 and PYK but did not repress PRPPS, and achieved robust *p*-CA production.

ST14207 exhibited both poor growth and low *p*-CA production. This strain combined the upregulation of TAL (Tyrosine Ammonia Lyase) and GCC2cm with the downregulation of PYK. The superior performance of ST14206, which also targets TAL and PYK but with a different TCA cycle enzyme (AKGDbm), points to the importance of the specific choice of TCA target in determining performance. Overexpression of GCC2cm likely disrupted TCA cycle flux, redox balance, or energy metabolism more severely than AKGDbm. Additionally, differences in enzyme kinetics, cofactor usage, or pathway integration between GCC2cm and AKGDbm may also have contributed to greater metabolic stress in ST14207.

These observations illustrate the complex trade-offs between growth and production that arise from specific genetic interventions, highlighting the need for data-driven strategies to guide design decisions. In this context, the predictive power of NOMAD increases as we incorporate data from different phenotypes and mutants^23^. For example, the data acquired on 10 mutant strains can be used to refine the kinetic models and improve future design steps. This data-driven foundation makes NOMAD well-suited for design–build–test–learn (DBTL) cycles, where experimental data generated in each round, including the reference strain and its derivatives, contribute to a deeper understanding of cellular physiology and are incorporated into model calibration. As model accuracy improves over successive cycles, so does their ability to identify rate-limiting steps and guide more effective interventions that further increase titers.

Moreover, NOMAD’s modeling principles are organism- and physiology-agnostic, and its predictions are contextualized through data integration across diverse conditions. This allows the framework to be readily adapted to different metabolic targets. Its capacity to incorporate heterogeneous datasets while preserving physiological proximity between strain designs ensures applicability and efficiency across a broad range of production objectives.

Our work highlights the potential of kinetic models to explore the design space systematically prior to experimental implementation. These models can be used for targeted low-throughput screening when high-throughput screening (HTS) is impossible – for example, when biosensors do not exist. They can also be combined with high-throughput screening first to identify combinatorial targets^29^ and then optimize the promoter strengths of the chosen targets. Despite their promise, the complexities and challenges of building kinetic models using traditional methods^15,63–66^ have meant that only a handful of large-scale models exist^67–70^. Recent advances in machine learning methods, such as RENAISSANCE^18^, ReKindle^20^, and iSCHRUNK^71–73^, as well as other approaches like KETCHUP^21^, have made it possible to parameterize and refine large-scale models rapidly. Coupled with the increasing availability of high-quality omics data, these developments will hasten the construction of large-scale models for a host of species^74^. These models can then be used to develop designs and test their viability in industrial settings by simulating their behavior in conditions that resemble real-world environments.

Overall, kinetic models offer a robust way to develop multi-target designs with a high likelihood of experimental success. This study paves the way toward a future in which libraries of large-scale kinetic models provide off-the-shelf solutions for various metabolic engineering applications, ultimately accelerating the development of microbial strains by minimizing the number of required design-build-test-learn cycles.

## Data and code availability

The raw data from the experiments, along with the modelling data, are provided in the Zenodo repository (https://doi.org/10.5281/zenodo.15432260) and the Supplementary Data file. The code used for generating the in-silico designs is available at https://github.com/EPFL-LCSB/NOMAD, in the ME-p-coumaric-acid directory.

## Acknowledgments

This work was supported by funding from the European Union’s Horizon 2020 research and innovation programme under grant agreement No. 814408 (SHIKIFACTORY100 project).

## Author contributions

**Bharath Narayanan:** Writing – original draft, Methodology, Conceptualization, Software, Formal analysis, Investigation, Visualization. **Wei Jiang**: Writing – original draft, Methodology, Conceptualization, Investigation, Formal Analysis, Visualization. **Shengbao Wang**: Writing – original draft, Methodology, Conceptualization, Investigation, Formal Analysis, Visualization. **Maria-Masid Barcon**: Methodology, Software. **Javier Sáez-Sáez**: Investigation, Analysis. **Viktor Hesselberg-Thomsen**: Investigation, Analysis. **Daniel Weilandt:** Methodology, Software. **Irina Borodina**: Resources, Conceptualization, Writing-Review and Editing, Supervision, Project administration, Funding acquisition. **Vassily Hatzimanikatis:** Resources, Formal analysis, Supervision, Funding acquisition. **Ljubisa Miskovic**: Conceptualization, Formal analysis, Writing-Review and Editing, Supervision, Project administration, Resources, Funding acquisition.

## Declaration of generative AI technologies in the writing process

During the preparation of this work, the author(s) used Grammarly and ChatGPT to improve language and text flow. After using this tool/service, the authors reviewed and edited the content as needed and take full responsibility for the content of the publication.

