## Supplementary material for "Kinetic-model-guided engineering of multiple *S. cerevisiae* strains improves *p*-coumaric acid production"

Supplementary information

### Supplementary Notes

**Supplementary Note I – sensitivity to choice of Vmax**

We built the nine kinetic models using steady-state data from ST10284 that was obtained during fed-batch fermentation. The final implementation of the designs was done in batch fermentation during which the strain has been observed to grow at roughly three times faster. In the absence of any other information on the behavior of the strain in batch fermentations, we accounted for this altered phenotype by multiplying all the maximal velocities in the kinetic model ($V_{max}s$) by three.

We wanted to analyze the sensitivity of our models to this approach of tripling the $V_{max}$s i.e. we wanted to know what the distribution of $V_{max}$s would be if we developed models from scratch with 3x the exofluxomic values while allowing all other variables to vary within 20% of their values under fed-batch conditions. We first sampled 100 steady-state profiles using the steady-state profile of the nine kinetic models as a scaffold, under the following conditions: (i) we tripled the exofluxomic values, (ii) we allowed all other fluxes to vary by 20% around 3x their original steady-state values, (iii) we allowed the concentrations to vary by 20% around their original steady-state values. We then sampled 20 sets of kinetic parameters around each of these 100 steady states while fixing the $K_{M}$s to their values from one of the nine kinetic models.

We then analysed the deviation of the $V_{max}$s for the 303 reactions in these 2,000 kinetic models around their fed-batch values. The histogram below shows the fold-change (in log-scale) for 606,000 (303 x 2000) $V_{max}$s (Supplementary Figure S2). The median value of the increase in $V_{max}$ was 2.9 (IQR: 2.3 – 3.4). 217 out of the 303 reactions had a median increase in $V_{max}$ that was within 25% of a 3x increase. Out of the remaining 86 reactions, 56 were transport reactions which we had assumed to be close to equilibrium and set as such. The other 30 reactions were also close to equilibrium; they had a median thermodynamic displacement of 0.96. The proximity of these 86 reactions to equilibrium meant that even slight changes in the ratios of the concentrations of their products and substrates lead to extremely large $V_{max}$s to ensure the same steady-state flux. This explains the large variations in the increase in their $V_{max}$s; we had permitted up to 20% change in the concentrations of the different metabolites while sampling. Notably, the fact that they are close to equilibrium precludes them from exerting control over the production pathway. This is because reactions close to equilibrium have higher elasticities and consequently have lower control coefficients as they are inversely proportional to the elasticities. Consequently, none of these reactions that had a median increase in $V_{max}$ that was outside the 25% margin formed part of any of the generated designs.

### Supplementary Tables

**Supplementary Table S1.** Strains used in this study.

| Strain No. | Genotype | Source |
| --- | --- | --- |
| CEN.PK113-7D | *MATa URA3 HIS3 LEU2 TRP1 MAL2-8c SUC2* | (Entian and Kötter, 2007) |
| CEN.PK113-13D | *CEN.PK113-7D-ΔURA3* | (Entian and Kötter, 2007) |
| ST10284 | *CEN.PK113-7D*-***ΔPDC5***-***ΔARO10***-***FjTAL***-*EcaroL*-***ΔGPP1***-***ScARO4^K229L^***-***ScARO7^G141S^***-*ScARO1*-*ScARO2*-***ScARO3***-*ScPHA2*-***AtPAL2***-*AtC4H*-*ScCYB5*-*AtCPR*-*MtPDH1*-***Bbxfpk*** | This study |
| ST14186 | ST10284*-ScAAT2*-*FjTAL*-*ScHXK2* | This study |
| ST14195 | ST10284*-PrPYK1*::*PrPRL18b-PrNDE1*::*PrREV1*-*FjTAL* | This study |
| ST14201 | ST10284*-PrPYK1*::*PrPRL18b*-*FjTAL* | This study |
| ST14204 | ST10284-*PrPYK1*::*PrPRL18b*-*PrPRS1::PrREV1*-*ScHXK2* | This study |
| ST14205 | ST10284*-ScAAT2-FjTAL-ScEMI2* | This study |
| ST14206 | ST10284*-PrPYK1::PrPRL18b::FjTAL-ScKGD1-ScKGD2-ScYMR31-ScLPD1* | This study |
| ST14207 | ST10284*-PrPYK1::PrPRL18b-FjTAL-ScGCV1-ScGCV2-ScGCV3-ScLPD1* | This study |
| ST14210 | ST10284-*PrPYK1::PrPRL18b-ScHXK2-ScADE6* | This study |
| ST14211 | ST10284*-PrPYK1::PrPRL18-EcaroB-PrPDC1::PrTEF1* | This study |
| ST14212 | ST10284*-EcaroB-EcaroD-PrPDC1::PrTEF1* | This study |
| ST14798 | CEN.PK113-13D-*PrPYK1*-*gfp-URA3* | This study |
| ST14799 | CEN.PK113-13D-*PrPDC1*-*gfp-URA3* | This study |
| ST14800 | CEN.PK113-13D-*PrTEF1*-*gfp-URA3* | This study |
| ST14801 | CEN.PK113-13D-*PrHHF1*-*gfp-URA3* | This study |
| ST14802 | CEN.PK113-13D-*PrRNR1*-*gfp-URA3* | This study |
| ST14803 | CEN.PK113-13D-*PrRPL18b*-*gfp-URA3* | This study |
| ST14804 | CEN.PK113-13D-*PrPAB1*-*gfp-URA3* | This study |
| ST14805 | CEN.PK113-13D-*PrRNR2*-*gfp-URA3* | This study |
| ST14806 | CEN.PK113-13D-*PrNDE1*-*gfp-URA3* | This study |
| ST14807 | CEN.PK113-13D-*PrPRS1*-*gfp-URA3* | This study |
| ST14808 | CEN.PK113-13D-*PrRAD27-gfp-URA3* | This study |
| ST14809 | CEN.PK113-13D-PrRAD27-gfp-URA3 | This study |

**Supplementary Table S2.** Chemical reaction targets suggested by NOMAD

| **Reactions (BIGG Models)** | **Chemical Reactions** |
| --- | --- |
| PYK | phosphoenolpyruvate + ADP + H+ → pyruvate + ATP |
| HEX1 | D-glucose + ATP → D-glucose 6-phosphate + ADP + H+ |
| PRPPS | D-ribose 5-phosphate + ATP ↔ 5-phospho-α-D-ribose 1-diphosphate + AMP + H+ |
| PRFGS | ATP + N2-formyl-N1-(5-phospho-β-D-ribosyl) glycinamide + L-glutamine + H2O → L-glutamate + ADP + 2-(formamido)-N1-(5-phospho-β-D-ribosyl) acetamidine + phosphate + H+ |
| AKGDbm | 2-oxoglutarate + coenzyme A + NAD+ → succinyl-CoA + CO2 + NADH |
| GCC2cm | glycine + a tetrahydrofolate + NAD+ → a 5,10-methylenetetrahydrofolate + ammonium + CO2 + NADH |
| ASPTA | 2-oxoglutarate + L-aspartate ⇄ oxaloacetate + L-glutamate |
| PYRDC | pyruvate + H+ → acetaldehyde + CO2 |
| DHQS | 3-deoxy-D-arabino-heptulosonate 7-phosphate → 3-dehydroquinate + phosphate |
| SUCOASm | succinate + ATP + coenzyme A ⇄ succinyl-CoA + ADP + phosphate |
| DHQTi | 3-dehydroquinate ⇄ 3-dehydroshikimate + H2O |
| NADH2_u6m | NADH + ubiquinone-6 → NAD+ + ubiquinol-6 |
| r_4235, EMI2 | D-hexose + ATP → D-hexose 6-phosphate + ADP + H+ |

**Supplementary Table S3.** Primers used in this study.

| **Primer** | **Sequence** | **Description** |
| --- | --- | --- |
| PR-32604 FWD | CGCAGAAAGTAATATCATGCGTCA | Check the homologous region of pCfB3041 |
| PR-32605 REV | TTTTTACGGTTCCTGGCCTTTT |  |
| PR-32614 FWD | TCGAGTCATGTAATTAGTTAATCGCG | Amplify pCfB3041 backbone for all gRNA plasmids construction used in this work |
| PR-32615 REV | AGCTCCAGCTTTTGTTCCCTATC |  |
| PR-32645 FWD | AGTGTCAACAACGTATCTACTTACGCTGATTGACAATCGG | Amplify *PrTEF1_EcaroB* from pCfB01955 |
| PR-32646 REV | CCTTTTCGGTTAGAGCGGATGCACACACCATAGCTTCAAAAT |  |
| PR-33619 FWD | CAGTTCGAGTTTATCATTATCAATACTGCC | Amplify *PrTDH3* as the repair template for upregulation |
| PR-33620 REV | TTTGTTTGTTTATGTGTGTTTATTCGAAAC |  |
| PR-33621 FWD | GTGTTGTTATCCGATACAACCGGATATTTT | Amplify *PrREV1* as the repair template for downregulation |
| PR-33622 REV | CGCTGGATATGCCTAGAAATGCAGT |  |
| PR-33623 FWD | CATGCGACTGGGTGAGCATATG | Amplify 500bp *PDC1* upstream HR containing 20bp overhangs to *PrREV1* |
| PR-33624 REV | TCCGGTTGTATCGGATAACAACACGGCAGCGGACAATATTATGCAATTATG |  |
| PR-33625 FWD | GCATTTCTAGGCATATCCAGCGATGTCTGAAATTACTTTGGGTAAATATTTG | Amplify 500bp *PDC1* downstream HR containing 20bp overhangs to *PrREV1* |
| PR-33626 REV | AAACCTAAGTAGACTGGTCTTTGGG |  |
| PR-33627 FWD | CCCTGGTCAAACTTCAGAACTAAG | Amplify 500bp *PYK1* upstream HR containing 20bp overhangs to *PrREV1* |
| PR-33628 REV | TCCGGTTGTATCGGATAACAACACCTTGGAAGACATCACAAGCATTC |  |
| PR-33629 FWD | GCATTTCTAGGCATATCCAGCGATGTCTAGATTAGAAAGATTGACCTC | Amplify 500bp *PYK1* downstream HR containing 20bp overhangs to *PrREV1* |
| PR-33630 REV | CCTTGACCTTCAAAGTCTTG |  |
| PR-33631 FWD | ATGAAACCCAAACTGTGGATATTATTTC | Amplify 500bp *NDE1* upstream HR containing 20bp overhangs to *PrREV1* |
| PR-33632 REV | TCCGGTTGTATCGGATAACAACACGAGAAGATCCCTGTCGCTTTC |  |
| PR-33633 FWD | GCATTTCTAGGCATATCCAGCGATGATTAGACAATCATTAATGAAAACAGTG | Amplify 500bp *NDE1* downstream HR containing 20bp overhangs to *PrREV1* |
| PR-33634 REV | CAATAGATTTCAATTCGATGGTACC |  |
| PR-33635 FWD | GACGTTTATGATTTGACTGGCTCC | Amplify 500bp *PRS1* upstream HR containing 20bp overhangs to *PrREV1* |
| PR-33636 REV | TCCGGTTGTATCGGATAACAACACATCGTGAAGGTTCGGTTATTGC |  |
| PR-33637 FWD | GCATTTCTAGGCATATCCAGCGATGCGTAAGTGTAAAATTTTTGTTGG | Amplify 500bp *PRS1* downstream HR containing 20bp overhangs to *PrREV1* |
| PR-33638 REV | CTCTAATCCATTTCGCTAAACTAGGG |  |
| PR-33639 FWD | CCGACTGATGTTTAGCTCAAGTTTG | Amplify 500bp *ADE6* upstream HR containing 20bp overhangs to *PrTDH3* |
| PR-33640 REV | TGATAATGATAAACTCGAACTGGGACAGCCAATTATTTAAATACGTATGTC |  |
| PR-33641 FWD | AATAAACACACATAAACAAACAAAATGACTGATTATATTTTGCCGGG | Amplify 500bp *ADE6* downstream HR containing 20bp overhangs to *PrTDH3* |
| PR-33642 REV | GTGTCATCCTATCGTAGACACACTTC |  |
| PR-33643 FWD | CTACAAATGAAAATCTAGGGCGC | Amplify 500bp *GCV1* upstream HR containing 20bp overhangs to *PrTDH3* |
| PR-33644 REV | TGATAATGATAAACTCGAACTGACTGTAATTGGATTTGTTTGGGTTC |  |
| PR-33645 FWD | AATAAACACACATAAACAAACAAAATGTCTATAATCAAAAAAATTGTGTTTAAGAGA | Amplify 500bp *GCV1* downstream HR containing 20bp overhangs to *PrTDH3* |
| PR-33646 REV | TGCAATGCCAATAGTGATCTACCT |  |
| PR-33651 FWD | GATATATGCGTTGACTCTGTTGC | Amplify 500bp *YMR31* upstream HR containing 20bp overhangs to *PrTDH3* |
| PR-33652 REV | TGATAATGATAAACTCGAACTGCCTGGGGAAGTACTCCTTAATG |  |
| PR-33653 FWD | AATAAACACACATAAACAAACAAAATGATTGCTACACCTATAAGATTAGCAAAG | Amplify 500bp *YMR31* downstream HR containing 20bp overhangs to *PrTDH3* |
| PR-33654 REV | CATAATATCATCACTCTTCTCTCTTCCTAAG |  |
| PR-33655 FWD | GCATCATCAAGAACGACGTTTAAG | Amplify 500bp *LPD1* upstream HR containing 20bp overhangs to *PrTDH3* |
| PR-33656 REV | TGATAATGATAAACTCGAACTGGCCGTTCGAGATGATAAAGTATTCT |  |
| PR-33657 FWD | AATAAACACACATAAACAAACAAAATGTTAAGAATCAGATCACTCCTAAATAATAAGC | Amplify 500bp *LPD1* downstream HR containing 20bp overhangs to *PrTDH3* |
| PR-33658 REV | ACATCTAGTATGTGGTCTTCCTTGACAG |  |
| PR-33659 FWD | ACGCTATTTTCTTCTTCCAGTTCATC | Amplify 500bp *LSC1* upstream HR containing 20bp overhangs to *PrTDH3* |
| PR-33660 REV | TGATAATGATAAACTCGAACTGCAAACAGAACAACGATTAGGGG |  |
| PR-33661 FWD | AATAAACACACATAAACAAACAAAATGTTAAGATCTACCGTTTCAAAAGC | Amplify 500bp *LSC1* downstream HR containing 20bp overhangs to *PrTDH3* |
| PR-33662 REV | CCAATTCTTACCTTTGTTGCTG |  |
| PR-33663 FWD | CATGCGTTCTACTACCAACAGAG | Amplify 500bp *LSC2* upstream HR containing 20bp overhangs to *PrTDH3* |
| PR-33664 REV | TGATAATGATAAACTCGAACTGAGAGAAAACGTCAAAGTGCTG |  |
| PR-33665 FWD | AATAAACACACATAAACAAACAAAATGTACTCAAGAAAATCCTTATCCCTAATTTC | Amplify 500bp *LSC2* downstream HR containing 20bp overhangs to *PrTDH3* |
| PR-33666 REV | CCACCTTGACTGGACGCAAT |  |
| PR-33667 FWD | CTGCATCAAGATCCCTCTAACTAC | Amplify 500bp *GCV3* upstream HR containing 20bp overhangs to *PrTDH3* |
| PR-33668 REV | TGATAATGATAAACTCGAACTGCCCTTTCCCATTAGATACTTGTTTG |  |
| PR-33669 FWD | AATAAACACACATAAACAAACAAAATGTTACGCACTACTAGACTATGGACC | Amplify 500bp *GCV3* downstream HR containing 20bp overhangs to *PrTDH3* |
| PR-33670 REV | CCAGTGTCTTTTCGTACTGTTCTAAG |  |
| PR-33671 FWD | ACGCTGGTAAAGTACAGCTACATTC | Amplify 500bp *HXK2* upstream HR containing 20bp overhangs to *PrTDH3* |
| PR-33672 REV | TGATAATGATAAACTCGAACTGCAGAAAGAAAAGAACGATTACCTAAAC |  |
| PR-33673 FWD | AATAAACACACATAAACAAACAAAATGGTTCATTTAGGTCCAAAAAAACC | Amplify 500bp *HXK2* downstream HR containing 20bp overhangs to *PrTDH3* |
| PR-33674 REV | GATTTTGTTTTGAGAAGCTGGG |  |
| PR-33675 FWD | CAAATAAGCAAAGAGAAACTTGAATGG | Amplify 500bp *AAT2* upstream HR containing 20bp overhangs to *PrTDH3* |
| PR-33676 REV | TGATAATGATAAACTCGAACTGGGTGACCATTAGTATTGACAATATGTACATC |  |
| PR-33677 FWD | AATAAACACACATAAACAAACAAAATGTCTGCCACTCTGTTCAATAAC | Amplify 500bp *AAT2* downstream HR containing 20bp overhangs to *PrTDH3* |
| PR-33678 REV | GAAAGCCGTTTAGGTCCAAAG |  |
| PR-33679 FWD | TCTTTTGCATCTTCCCTTGGGAG | Amplify 500bp *GCV2* upstream HR containing 20bp overhangs to *PrTDH3* |
| PR-33680 REV | TGATAATGATAAACTCGAACTGGATGGTTAGATCGCGCAGCAATC |  |
| PR-33681 FWD | AATAAACACACATAAACAAACAAAATGCTTAGGACAAGAGTGACTGC | Amplify 500bp *GCV2* downstream HR containing 20bp overhangs to *PrTDH3* |
| PR-33682 REV | GTATACCATTCTGGACTTTCTAGCAG |  |
| PR-33683 FWD | TACAGTAGTAGTCTCAGCACCCC | Amplify 500bp *EMI2* upstream HR containing 20bp overhangs to *PrTDH3* |
| PR-33684 REV | TGATAATGATAAACTCGAACTGGATCTTCTTCCTCTGTCTTCCA |  |
| PR-33685 FWD | AATAAACACACATAAACAAACAAAATGTCATTTGAAAATTTACATAAAGTCAATG | Amplify 500bp *EMI2* downstream HR containing 20bp overhangs to *PrTDH3* |
| PR-33686 REV | GATATGAGAAGGTAAATCCCATTTTC |  |
| PR-33687 FWD | GTGGGACATATTCGAACGTCTC | Amplify 500bp *KGD2* upstream HR containing 20bp overhangs to *PrTDH3* |
| PR-33688 REV | TGATAATGATAAACTCGAACTGCTGATATATACCGTTAAAGATAGCTGCCT |  |
| PR-33689 FWD | AATAAACACACATAAACAAACAAAATGCTTTCCAGAGCGACGC | Amplify 500bp *KGD2* downstream HR containing 20bp overhangs to *PrTDH3* |
| PR-33690 REV | GTTCGGTAGGCTCTGGCTTAGATTC |  |
| PR-33691 FWD | GACTATATCCCTTTCCCCTCTAATACTC | Amplify 500bp *KGD1* upstream HR containing 20bp overhangs to *PrTDH3* |
| PR-33692 REV | TGATAATGATAAACTCGAACTGGAAATTAGGAAGAAGTATGATGCTGAA |  |
| PR-33693 FWD | AATAAACACACATAAACAAACAAAATGCTAAGGTTCGTGTCTTCG | Amplify 500bp *KGD1* downstream HR containing 20bp overhangs to *PrTDH3* |
| PR-33694 REV | CCGTAGTAGTCTAGAGTCAATTCCG |  |
| PR-33695 FWD | GCCAGATTAAAATTCACGAACTCTTC | Yeast colony PCR forward primer for genes regulated by *PrREV1* |
| PR-33696 REV | CCAGGGCTGTAACTCTTTTAGTACCAC | Yeast colony PCR reverse primer binds *PRS1*; pair with PR-33695 |
| PR-33697 REV | GCTTCAGCTTCATAGTAATGGACTTC | Yeast colony PCR reverse primer binds *NED1*; pair with PR-33695 |
| PR-33698 REV | GCAAATCGACATCGGTACCTG | Yeast colony PCR reverse primer binds *PYK1*; pair with PR-33695 |
| PR-33699 REV | CGTTTGGCTTCAAAGACATGTC | Yeast colony PCR reverse primer binds *PDC1*; pair with PR-33695 |
| PR-33700 FWD | CTTCACCAACCATCAGTTCATAGGTC | Yeast colony PCR forward primer for genes regulated by *PrTDH3* FWD For Colony PCR |
| PR-33701 REV | CTTGCAAACCTTGGCAGGATAC | Yeast colony PCR reverse primer binds *KGD1*; pair with PR-33700 |
| PR-33702 REV | CCGACCGTGTCTTCAATCTTG | Yeast colony PCR reverse primer binds *EMI2*; pair with PR-33700 |
| PR-33703 REV | CAACATTGGAACAACATCGTGG | Yeast colony PCR reverse primer binds *HXK2*; pair with PR-33700 |
| PR-33704 REV | CGAAAGAGCACCTGTTGAAGAGAC | Yeast colony PCR reverse primer binds *LPD1*; pair with PR-33700 |
| PR-33705 REV | GCTTTCTCATTTGCAATGCTTATCTC | Yeast colony PCR reverse primer binds *GCV1*; pair with PR-33700 |
| PR-33706 REV | GCCAAGGATATTAAAGGCTACGGT | Yeast colony PCR reverse primer binds *YMR31*; pair with PR-33700 |
| PR-33707 REV | GTACGTATAAGCTTTGTTCAGTCATCATG | Yeast colony PCR reverse primer binds *GCV3*; pair with PR-33700 |
| PR-33709 REV | GTTTGCTGAACAGCTTCGTATGTC | Yeast colony PCR reverse primer binds *LSC1*; pair with PR-33700 |
| PR-33710 REV | GAAAATTTCTTTATAGCATCTGGGGTTC | Yeast colony PCR reverse primer binds *LSC2*; pair with PR-33700 |
| PR-33711 REV | CTCATCGTATTTGGTGGTTCGG | Yeast colony PCR reverse primer binds *ADE6*; pair with PR-33700 |
| PR-33712 REV | CGATTTGAACCCATTGTTCACTAG | Yeast colony PCR reverse primer binds *AAT2*; pair with PR-33700 |
| PR-33713 REV | CCGTTTGAAAGTTTAATAGCGCTTC | Yeast colony PCR reverse primer binds *GCV2*; pair with PR-33700 |
| PR-33714 REV | CTTCTTTCTTTGGAGCGGCTTC | Yeast colony PCR reverse primer binds *KGD2*; pair with PR-33700 |
| PR-33717 FWD | AGTGTCAACAACGTATCTACGCACACACCATAGCTTCAAA | Amplify *PrTEF1* for constructing single overexpression plasmids through Gibson assembly |
| PR-33718 REV | TTGTAATTAAAACTTAGATTAGATTGCTATG |  |
| PR-33719 FWD | AATCTAAGTTTTAATTACAAATGTCTGCCACTCTGTTCAA | Amplify *AAT2* from yeast genome for constructing single overexpression plasmids through Gibson assembly. |
| PR-33720 REV | CCTTTTCGGTTAGAGCGGATTTACAATTTAGCTTCAGTAGCATAGAAG |  |
| PR-33723 FWD | AATCTAAGTTTTAATTACAAATGGTTCATTTAGGTCCAAA | Amplify *HXK2* from yeast genome for constructing single overexpression plasmids through Gibson assembly. |
| PR-33724 REV | CCTTTTCGGTTAGAGCGGATTTAAGTTTAAGCACCGATGA |  |
| PR-33725 FWD | AATCTAAGTTTTAATTACAAATGTCATTTGAAAATTTACATAAAGTCAATG | Amplify *EMI2* from yeast genome for constructing single overexpression plasmids through Gibson assembly. |
| PR-33726 REV | CCTTTTCGGTTAGAGCGGATTTATGCCACCAGAGCACACA |  |
| PR-33727 FWD | AATCTAAGTTTTAATTACAAATGTTAAGATCTACCGTTTCAA | Amplify *LSC1* from yeast genome for constructing single overexpression plasmids through Gibson assembly. |
| PR-33728 REV | CCTTTTCGGTTAGAGCGGATTCATTTAAATTTGGCAAATT |  |
| PR-33729 FWD | AATCTAAGTTTTAATTACAAATGTACTCAAGAAAATCCTTATCC | Amplify *LSC2* from yeast genome for constructing single overexpression plasmids through Gibson assembly. |
| PR-33730 REV | CCTTTTCGGTTAGAGCGGATTTAATTTTGGGTCAATTCAA |  |
| PR-33731 FWD | TTGTTTTATATTTGTTGTAAAAAGTAG | Amplify *PrTEF1* and *PrPGK1* for constructing double overexpression plasmids through Gibson assembly |
| PR-33732 REV | TTGTAATTAAAACTTAGATTAGATTGC |  |
| PR-33737 FWD | AGTGTCAACAACGTATCTACTCAACAATGAATAGCTTTATCA | Amplify *LPD1* from yeast genome for constructing double overexpression plasmids through Gibson assembly (link with *GCV3*) |
| PR-33738 REV | TTACAACAAATATAAAACAAATGTTAAGAATCAGATCACTCC |  |
| PR-33739 FWD | AATCTAAGTTTTAATTACAAATGTTACGCACTACTAGACTATGG | Amplify *GCV3* from yeast genome for constructing double overexpression plasmids through Gibson assembly |
| PR-33740 REV | CCTTTTCGGTTAGAGCGGATACCAGTGTCTTTTCGTACTG |  |
| PR-33745 FWD | AGTGTCAACAACGTATCTACTCAACAATGAATAGCTTTATCA | Amplify *LPD1* from yeast genome for constructing double overexpression plasmids through Gibson assembly (link with *YMR31*) |
| PR-33746 REV | TTACAACAAATATAAAACAAATGTTAAGAATCAGATCACTCC |  |
| PR-33747 FWD | AATCTAAGTTTTAATTACAAATGATTGCTACACCTATAAGATTAGCAAAGAGTG | Amplify *YMR31* from yeast genome for constructing double overexpression plasmids through Gibson assembly |
| PR-33748 REV | CCTTTTCGGTTAGAGCGGATTCACCATGCACCACCGCTAT |  |
| PR-33773 FWD | GGTCTCAAACGAAAATGAAGGCCAAATCAAGGC | Amplify the region 1kb upstream of *PDC1* for fusion with *SfGFP* using Golden Gate assembly on pYTK096 |
| PR-33774 REV | GGTCTCACATATTTGATTGATTTGACTGTGTTATTTTGC |  |
| PR-33775 FWD | GGTCTCAAACGACTTGAGATGTGTGTCAATGCTAGTATTTTG | Amplify the region 1kb upstream of *PYK1* for fusion with *SfGFP* using Golden Gate assembly on pYTK096 |
| PR-33776 REV | GGTCTCACATATGTGATGATGTTTTATTTGTTTTGATTGG |  |
| PR-33777 FWD | GGTCTCAAACGCTACAACGTTGAAAACCAACAATATGTTG | Amplify the region 1kb upstream of *PRS1* for fusion with *SfGFP* using Golden Gate assembly on pYTK096 |
| PR-33778 REV | GGTCTCACATATTTTATTTCTCTTCAAATTAAGACTATTAAACGGTAG |  |
| PR-33779 FWD | GGTCTCAAACGGAAACCCAAACTGTGGATATTATTTC | Amplify the region 1kb upstream of *NDE1* for fusion with *SfGFP* using Golden Gate assembly on pYTK096 |
| PR-33780 REV | GGTCTCACATATATTATTGGTTAATTTTTTATTTGCTCTAATAAGTC |  |
| PR-33781 FWD | AATCTAAGTTTTAATTACAAATGACTGATTATATTTTGCCGGGTC | Amplify *ADE6* from yeast genome for constructing single overexpression plasmids through Gibson assembly. |
| PR-33782 REV | CCTTTTCGGTTAGAGCGGATTCAACCGACCCATCTTCTGG |  |
| PR-33783 REV | CTGGTGGTAAGGCCCCAAGACTTGGAAGACATCACAAGCATTC | Amplify 500 bp upstream of *PYK1* with 20bp overhangs to *PrHHF1,* pair with PR-33627 |
| PR-33784 FWD | AAACAAGCAACAAATATAATATAGTAAAATATGTCTAGATTAGAAAGATTGACCTC | Amplify 500 bp downstream of *PYK1* with 20bp overhangs to *PrHHF1,* pair with PR-33630 |
| PR-33785 REV | CTGGTGGTAAGGCCCCAAGAGGCAGCGGACAATATTATGCAATTATG | Amplify 500 bp upstream of *PDC1* with 20bp overhangs to *PrHHF1,* pair with PR-33623 |
| PR-33786 FWD | AAACAAGCAACAAATATAATATAGTAAAATATGTCTGAAATTACTTTGGGTAAATATTTG | Amplify 500 bp downstream of *PDC1* with 20bp overhangs to *PrHHF1,* pair with PR-33626 |
| PR-33787 REV | CCATATTTGCAATTTCACAAGGGAGAAGATCCCTGTCGCTTTC | Amplify 500 bp upstream of *NDE1* with 20bp overhangs to *PrRAD27,* pair with PR-33631 |
| PR-33788 FWD | CGGACACCGGAAGAAAAAATATGATTAGACAATCATTAATGAAAACAGTG | Amplify 500 bp downstream of *NDE1* with 20bp overhangs to *PrRAD27,* pair with PR-33634 |
| PR-33789 REV | CCATATTTGCAATTTCACAAGGGACGTTTATGATTTGACTGGCTCC | Amplify 500 bp upstream of *PRS1* with 20bp overhangs to *PrRAD27,* pair with PR-33635 |
| PR-33790 FWD | CGGACACCGGAAGAAAAAATATCGTGAAGGTTCGGTTATTGC | Amplify 500 bp downstream of *PRS1* with 20bp overhangs to *PrRAD27,* pair with PR-33638 |
| PR-33791 FWD | TCTTGGGGCCTTACCACCAGT | Amplify *PrHFF1* from pYTK015 |
| PR-33792 REV | ATTTTACTATATTATATTTGTTGCTTGTTTTTGTTTGTT |  |
| PR-33793 FWD | CCTTGTGAAATTGCAAATATGGTG | Amplify *PrRAD27* from pYTK025 |
| PR-33794 REV | ATTTTTTCTTCCGGTGTCCGTT |  |
| PR-33795 FWD | AAGAGGATGTCCAATATTTTTTTTAAGGA | Amplify *PrPRL18b* from pYTK017 |
| PR-33796 REV | TTTGTTTTTTGTTTTCTTCTAATTGATTTTTT |  |
| PR-33805 FWD | CCTTGCCAACAGGGAGTTCTT | Amplify *PrTEF1* from pCfB01955 |
| PR-33806 REV | TTTGTAATTAAAACTTAGATTAGATTGCTATGCTTTC |  |
| PR-33807 REV | AAGAACTCCCTGTTGGCAAGGGGCAGCGGACAATATTATGCAATTATG | Amplify 500 bp upstream of *PDC1* with 20bp overhangs to *PrTEF1,* pair with PR-33623 |
| PR-33808 FWD | TTAAAACTTAGATTAGATTGCTATGCTTTCATGTCTGAAATTACTTTGGGTAAATATTTG | Amplify 500 bp downstream of *PDC1* with 20bp overhangs to *PrTEF1,* pair with PR-33626 |
| PR-34049 FWD | AGTGCAGGUAAAACAATGAATACGATTAATGAATATCTTTCACTTG | Amplify FjTAL with overhangs for USER cloning |
| PR-34050 REV | CGTGCGAUTTAATTATTGATCAAGTGATCTTTCACTTTC |  |
| PR-33797 FWD | CGTGCGAUTTAGGATTGTTGGAAAACATCTTTC | Amplify *KGD1* from yeast genome for constructing double overexpression plasmids through USER cloning (link with *KGD2*) |
| PR-33798 REV | AGTGCAGGUAAAACAATGCTAAGGTTCGTGTCTTCG |  |
| PR-33799 FWD | ATCTGTCAUAAAACAATGCTTTCCAGAGCGACGC | Amplify *KGD2* from yeast genome for constructing double overexpression plasmids through USER cloning (link with *KGD1*) |
| PR-33800 REV | CACGCGAUTCACCATAACAACATTTTTCTAGGGTCTT |  |
| PR-33801 FWD | CGTGCGAUTTACTGCTTGTAGTAATGTGTGGGC | Amplify *GCV1* from yeast genome for constructing double overexpression plasmids through USER cloning (link with *GCV2*) |
| PR-33802 REV | AGTGCAGGUAAAACAATGTCTATAATCAAAAAAATTGTGTTTAAGAGATTC |  |
| PR-33803 FWD | ATCTGTCAUAAAACAATGCTTAGGACAAGAGTGACTGC | Amplify *GCV2* from yeast genome for constructing double overexpression plasmids through USER cloning (link with *GCV1*) |
| PR-33804 REV | CACGCGAUTCATTCAGTTTCGTTCGCAA |  |

**Supplementary Table S4.** Biobricks used in this study.

| **ID** | **Name** | **Description** |
| --- | --- | --- |
| BB7435 | *EcAroB-EcAroD* | Codon optimized *EcAroB-EcAroD* for *S. cerevisiae* |
| BB7436 | *PrTEF1*->*EcAroB* | *EcAroD* regulated by *PrTEF1* |
| BB7443 | Upstream HR *NDE1* | 500bp upstream HR of *NDE1* with overhangs to *PrREV1* |
| BB7444 | Downstream HR *NDE1* | 500bp downstream HR of *NDE1* with overhangs to *PrREV1* |
| BB7445 | *PrNDE1::PrREV1* | 500bp repair template for *PrNDE1* swapped by *PrREV1* |
| BB7446 | Upstream HR *PRS1* | 500bp upstream HR of *PRS1* with overhangs to *PrREV1* |
| BB7447 | Downstream HR *PRS1* | 500bp downstream HR of *PRS1* with overhangs to *PrREV1* |
| BB7448 | *PrPRS1::PrREV1* | 500bp repair template for *PrPRS1* swapped by *PrREV1* |
| BB7449 | *AAT2* | Amplified from genome for the overexpression |
| BB7450 | *HXK2* | Amplified from genome for the overexpression |
| BB7451 | *EMI2* | Amplified from genome for the overexpression |
| BB7452 | *LSC1* | Amplified from genome for the overexpression |
| BB7453 | *LSC2* | Amplified from genome for the overexpression |
| BB7454 | *PrPGK1-PrTEF1* | Double primer for the double overexpression plasmid |
| BB7457 | *LPD1* | Amplified from genome for the double overexpression |
| BB7460 | *GCV3* | Amplified from genome for the overexpression, link with *LPD1* |
| BB7461 | *YMR31* | Amplified from genome for the overexpression, link with *LPD1* |
| BB7462 | Upstream HR *PYK1* | 500bp upstream HR of *PYK1* with overhangs to *PrPRL18b* |
| BB7463 | Downstream HR *PYK1* | 500bp downstream HR of *PYK1* with overhangs to *PrPRL18b* |
| BB7464 | *PrPYK1::PrPRL18b* | 500bp repair template for *PrPYK1* swapped by *PrPRL18b* |
| BB7474 | *PDC1* -1kb | Amplified from genome, with overhangs to GFP |
| BB7475 | *PYK1* -1kb | Amplified from genome, with overhangs to GFP |
| BB7476 | *PRS1* -1kb | Amplified from genome, with overhangs to GFP |
| BB7477 | *NDE1* -1kb | Amplified from genome, with overhangs to GFP |
| BB7478 | *ADE6* | Amplified from genome for the overexpression |
| BB7479 | *KGD1* USER | Amplified from genome for the double overexpression, with USER Cloning overhangs to *KGD2* |
| BB7480 | *KGD2* USER | Amplified from genome for the double overexpression, with USER Cloning overhangs to *KGD1* |
| BB7481 | *GCV1* USER | Amplified from genome for the double overexpression, with USER Cloning overhangs to *GCV2* |
| BB7482 | *GCV2* USER | Amplified from genome for the double overexpression, with USER Cloning overhangs to *GCV1* |
| BB7484 | Upstream HR *PDC1* | 500bp upstream HR of *PDC1* with overhangs to *PrTEF1* |
| BB7485 | Downstream HR *PDC1* | 500bp upstream HR of *PDC1* with overhangs to *PrTEF1* |
| BB7486 | *PrPDC1::PrTEF1* | 500bp repair template for *PrPDC1* swapped by *PrTEF1* |
| BB7487 | *PrHHF1* | Amplified from *pYTK015, with* overhangs to GFP |
| BB7488 | *PrRAD27* | Amplified from *pYTK025, with* overhangs to GFP |
| BB7489 | *PrPRL18b* | Amplified from *pYTK017, with* overhangs to GFP |
| BB7490 | *PrTEF1* | Amplified from *pCfB01955, with* overhangs to GFP |
| BB7491 | *FjTAL USER* | Amplified from DNA string for the overexpression |
| BB7492 | *gRNA_GCV2* | IDT DNA string |
| BB7493 | *gRNA_KGD2* | IDT DNA string |
| BB7494 | *gRNA_PYK1* | IDT DNA string |
| BB7495 | *gRNA_AAT2* | IDT DNA string |
| BB7496 | *gRNA_ADE6* | IDT DNA string |
| BB7497 | *gRNA_GCV3* | IDT DNA string |
| BB7498 | *gRNA_KGD1* | IDT DNA string |
| BB7499 | *gRNA_PRS1* | IDT DNA string |
| BB7500 | *gRNA_YMR31* | IDT DNA string |
| BB7501 | *gRNA_NDE1* | IDT DNA string |
| BB7502 | *gRNA_LSC1* | IDT DNA string |
| BB7503 | *gRNA_LPD1* | IDT DNA string |
| BB7504 | *gRNA_HXK2* | IDT DNA string |
| BB7505 | *gRNA_GCV1* | IDT DNA string |
| BB7506 | *gRNA_EMI2* | IDT DNA string |
| BB7507 | *gRNA_PDC1* | IDT DNA string |
| BB7507 | *gRNA_PDC1* | IDT DNA string |

**Supplementary Table S5.** Plasmids used in this study.

| **Name** | **Description** | **Parent plasmid** | **Biobricks** | **Reference** |
| --- | --- | --- | --- | --- |
| pCfB2312 | *PrTEF1-Cas9-TCYC1_KanMX* | See ref. | See ref. | (Stovicek et al., 2015) |
| pCfB3034 | EasyClone-MarkerFree backbone vector for the addition of 1 or 2 genes and promoters for integration into site X-3 | See ref. | See ref. | (Jessop-Fabre et al., 2016) |
| pCfB3041 | EasyClone-MarkerFree guiding RNA vector to direct Cas9 to cut at site X-3 | See ref. | See ref. | (Jessop-Fabre et al., 2016) |
| pCfB3044 | EasyClone-MarkerFree guiding RNA vector to direct Cas9 to cut at site XI-2 | See ref. | See ref. | (Jessop-Fabre et al., 2016) |
| pCfB3048 | EasyClone-MarkerFree guiding RNA vector to direct Cas9 to cut at site XII-2 | See ref. | See ref. | (Jessop-Fabre et al., 2016) |
| pCfB3051 | EasyClone-MarkerFree guiding RNA vector to direct Cas9 to cut at sites X-3, XI-2, and XII-2 | See ref. | See ref. | (Jessop-Fabre et al., 2016) |
| pCfB12642 | *IntX-3_TADH1<-EcAroB<-PrPGK1-PrTEF1->EcAroD->TCYC1* | pCfB3034 | BB7435  BB7454 | This study |
| pCfB12643 | *IntX-3_PrTEF1->EcAroB->TCYC1* | pCfB3034 | BB7436 | This study |
| pCfB12655 | *gRNA_PYK1_PDC1_NAT* | pCfB3041 | BB7494  BB7507 | This study |
| pCfB12656 | *gRNA_ PDC1_NAT* | pCfB3041 | BB7507 | This study |
| pCfB12658 | *IntXII_2_pTEF1_LSC2* | pCfB3048 | BB7453 | This study |
| pCfB12659 | *IntXII_2_pTEF1_EMI2* | pCfB3048 | BB7451 | This study |
| pCfB12660 | *IntX_3_pTEF1_AAT2* | pCfB3034 | BB7449 | This study |
| pCfB12661 | *IntXI_2_pTEF1_LSC1* | pCfB3044 | BB7452 | This study |
| pCfB12663 | *IntXII_2_pTEF1_HXK2* | pCfB3048 | BB7450 | This study |
| pCfB12664 | *IntXII-2_TADH1<-LPD1<-PrPGK1-PrTEF1->GCV3->TCYC1* | pCfB3048 | BB7454 BB7457 BB7460 | This study |
| pCfB12665 | *IntXII-2_TADH1<-LPD1<-PrPGK1-PrTEF1->YMR31->TCYC1* | pCfB3048 | BB7454 BB7457 BB7461 | This study |
| pCfB12666 | *PrTEF1->SfGFP->TENO1* | pYTK096 | BB7490 | (Lee et al., 2015) |
| pCfB12667 | *PrHHF1->SfGFP->TENO1* | pYTK096 | BB7487 | (Lee et al., 2015) |
| pCfB12668 | *PrPRL18b->SfGFP->TENO1* | pYTK096 | BB7489 | (Lee et al., 2015) |
| pCfB12672 | *PrRAD27->SfGFP->TENO1* | pYTK096 | BB7488 | (Lee et al., 2015) |
| pCfB12673 | *PrPYK1->SfGFP->TENO1* | pYTK096 | BB7474 | (Lee et al., 2015) |
| pCfB12674 | *PrPRS1->SfGFP->TENO1* | pYTK096 | BB7475 | (Lee et al., 2015) |
| pCfB12675 | *PrPDC1->SfGFP->TENO1* | pYTK096 | BB7476 | (Lee et al., 2015) |
| pCfB12676 | *PrNDE1->SfGFP->TENO1* | pYTK096 | BB7477 | (Lee et al., 2015) |
| pCfB12677 | *gRNA_ PYK1_NAT* | pCfB3041 | BB7494 | This study |
| pCfB12678 | *gRNA_ PYK1_PRS1_NAT* | pCfB3041 | BB7494 BB7499 | This study |
| pCfB12679 | *IntX_3_PrTEF1_ADE6* | pCfB3034 | BB7478 | This study |
| pCfB12680 | *IntX_3_ TADH1<-GCV1<-PrPGK1-PrTEF1->GCV2->TCYC1_USER* | pCfB3034 | BB7454 BB7481 BB7482 | This study |
| pCfB12681 | *IntX_3_ TADH1<-KGD1<-PrPGK1-PrTEF1->KGD2->TCYC1_USER* | pCfB3034 | BB7454 BB7479 BB7480 | This study |
| pCfB12682 | *gRNA_ PYK1_NDE1_NAT* | pCfB3041 | BB7494 BB7501 | This study |
| pCfB12683 | *gRNA_X-3_XI-2_NAT* | pCfB3041 | N/A | This study |
| pCfB12684 | *gRNA_ X-3_XII-2_NAT* | pCfB3041 | N/A | This study |
| pCfB12685 | *gRNA_ XI-2_XII-2_NAT* | pCfB3041 | N/A | This study |

**Supplementary Table S6.** Steady-state parameters of ST10284 obtained during fed-batch fermentation. We built the kinetic models using this phenotype and then tripled all $V_{max}s$ to approximate the behavior in a batch setting

| **Parameter** | **ST10284** |
| --- | --- |
| $\mu(h^{-1})$ | $0.097\pm0.004$ |
| $q_{glc}$ (mmol / gDW / h) | $-1.142\pm0.035$ |
| $q_{pca}$ (mmol / gDW / h) | $0.085\pm0.004$ |
| $q_{etoh}$ (mmol / gDW / h) | $0.007\pm0.002$ |

### Supplementary Figures

**Supplementary Figure S1:** Genetic engineering to enhance *p*-CA production in ST10284. DAHP synthase (*ARO3*), L-tyrosine-feedback-insensitive DAHP synthase (*ARO4^K229L^*), Pentafunctional aromatic protein (*ARO1*), Chorismate synthase (*ARO2*), L-tyrosine-feedback-insensitive chorismate mutase (*ARO7^G141S^*), Prephenate dehydratase (*PHA2), Arabidopsis thaliana* phenylalanine ammonia lyase (*AtPAL2*), Cinnamic acid hydroxylase (*AtC4H*), P450 reductase (*AtATR2*),Cytochrome b5 (*CYB5*), *Flavobacterium johnsoniae* tyrosine ammonia lyase (*FjTAL*), *E. coli* shikimate kinase (*EcAroL*), *Medicago truncatula* L-tyrosine prephenate dehydrogenase (*MtPDH1*), *Bifidobacterium breve* phosphoketolase (*Bbxfpk*), Phenylpyruvate decarboxylase (*ARO10*), pyruvate decarboxylase(*PDC10*), Glycerol-3-Phosphate Phosphatase (*GPP1*), Glc glucose, G6P glucose-6-phosphate, F6P fructose-6-phosphate, PEP phosphoenolpyruvate, E4P erythrose-4-phosphate, Pyr pyruvate, Ac-P acetyl-phosphate, Ac-CoA acetyl-CoA, TCA tricarboxylic acid cycle, DAHP 3-deoxy-D-arabino-2-heptulosonic acid 7-phosphate, EPSP 5-enolpyruvyl-shikimate-3-phosphate, CHA chorismic acid, PPA prephenate, PPY phenylpyruvate, HPP para-hydroxy-phenylpyruvate, L-Phe L-phenylalanine, L-Tyr L-tyrosine, CA cinnamic acid, p-CA p-coumaric acid. Native gene overexpression (Green), heterologous gene overexpression (Blue), Gene deletion (Red), feedback resistant variants of native genes(Gray).

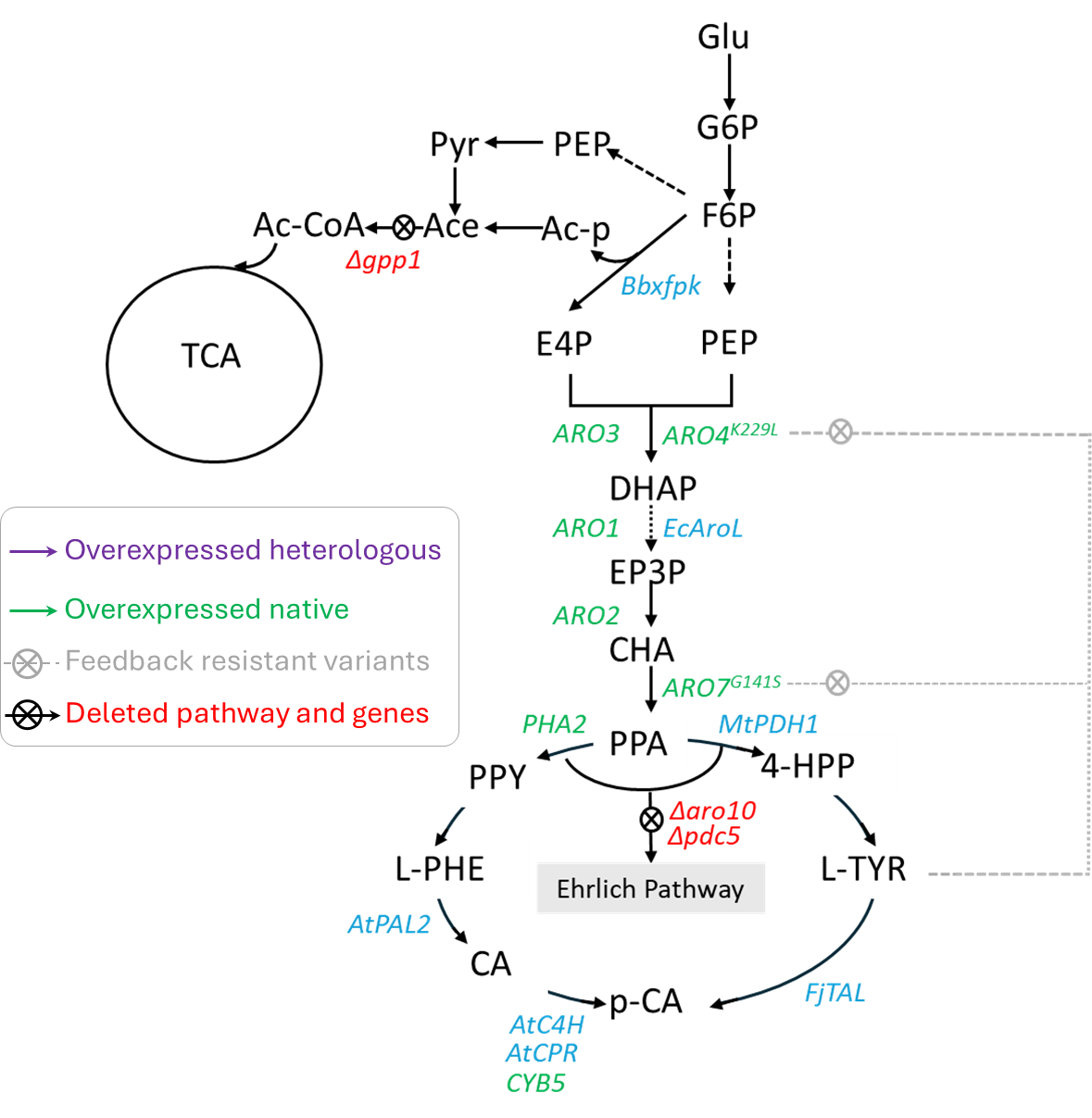

**Supplementary Figure S2:** Distribution of the increase in 303 Vmaxs across the 2,000 newly generated kinetic models (total of 606,000 Vmaxs). The x-axis is in log scale. The vertical bars represent a 25% margin on a 3-fold increase in Vmax when compared to the original kinetic model developed using fed-batch data.

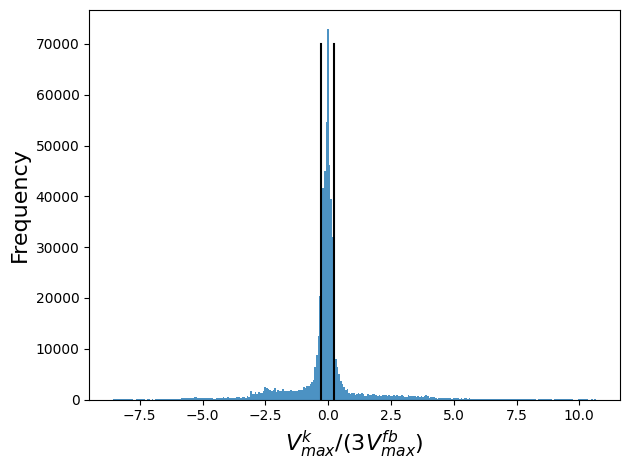

**Supplementary Figure S3:** Extracellular glucose concentrations in batch fermentation simulations for the 39 combinations of design and model (out of 90) that demonstrated higher titers than the 9 kinetic models representative of ST10284. This figure complements Figures 2B and 2C showing the biomass and *p*-CA titers. The dynamics of glucose uptake of the engineering strains (blue) was identical to those of the reference strain (orange).

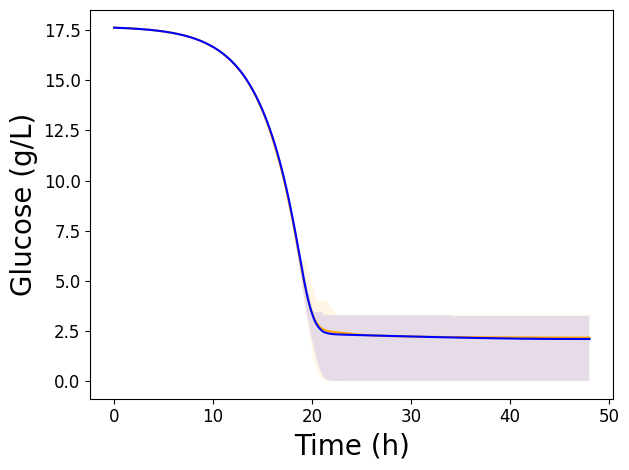

**Supplementary Figure S4:** Extracellular glucose concentrations in batch fermentation experiments for each of the 10 engineering strains and ST10284. For all the strains, glucose was depleted within 24 hours

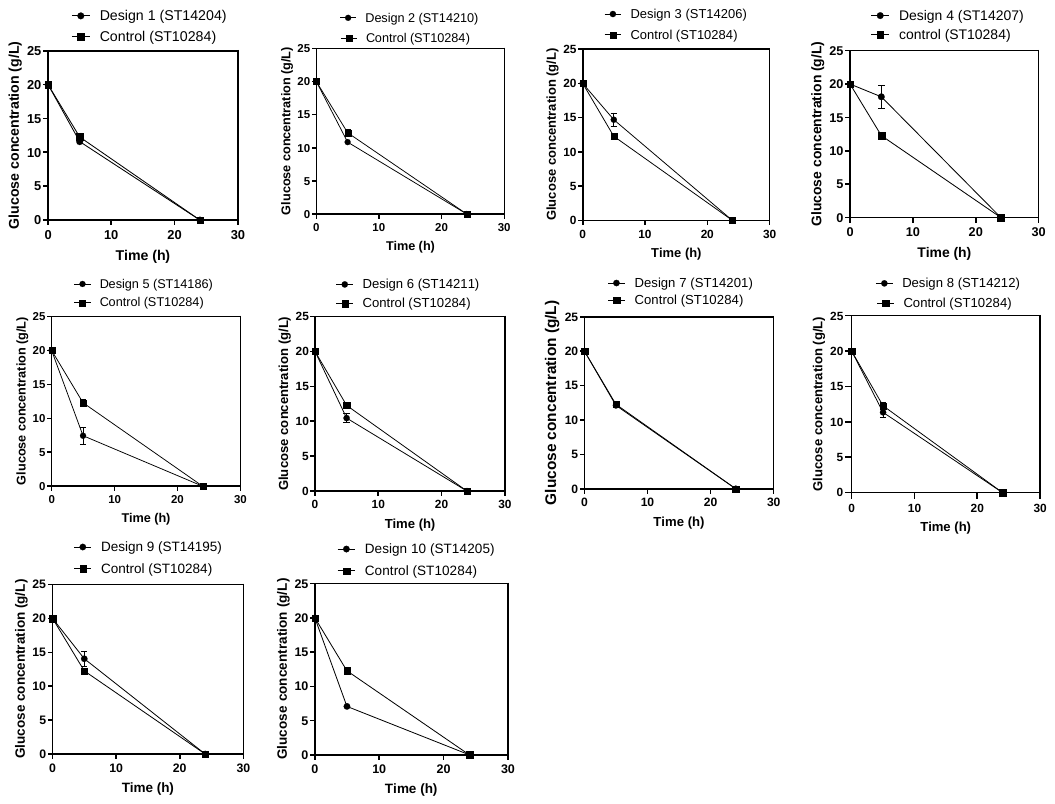

**Supplementary Figure S5:** Extracellular ethanol concentrations in batch fermentation experiments for the 10 engineering strains and ST10284. The strains secrete ethanol while growing on glucose. Eight out of the ten strains, except Design 1 and Design 4, completely consume the secreted ethanol to fuel further growth and p-CA production.

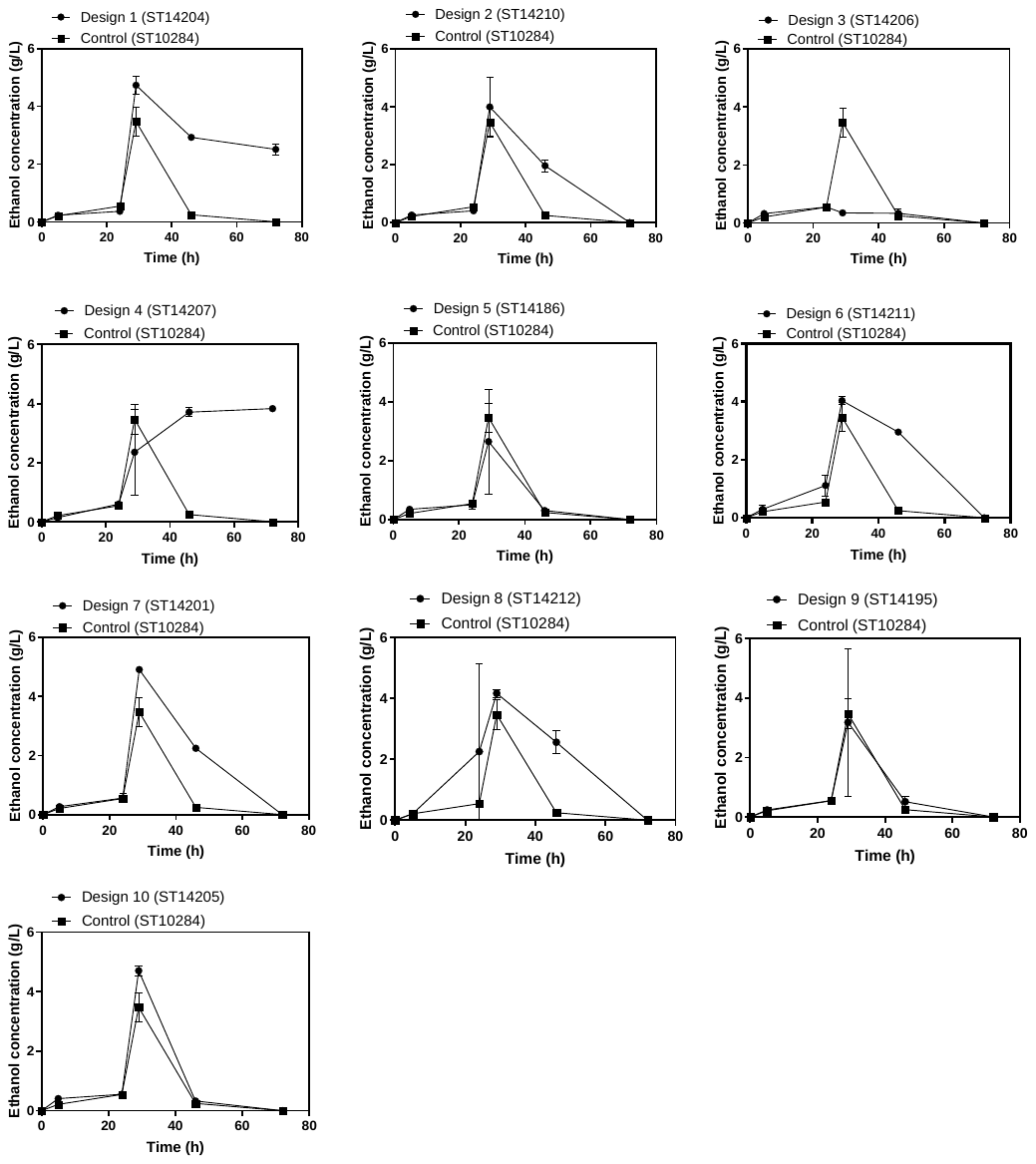

**Supplementary Figure S6:** Growth and *p*-CA titers from the batch fermentation experiments for the 10 engineering strains and the reference strain (ST10284).

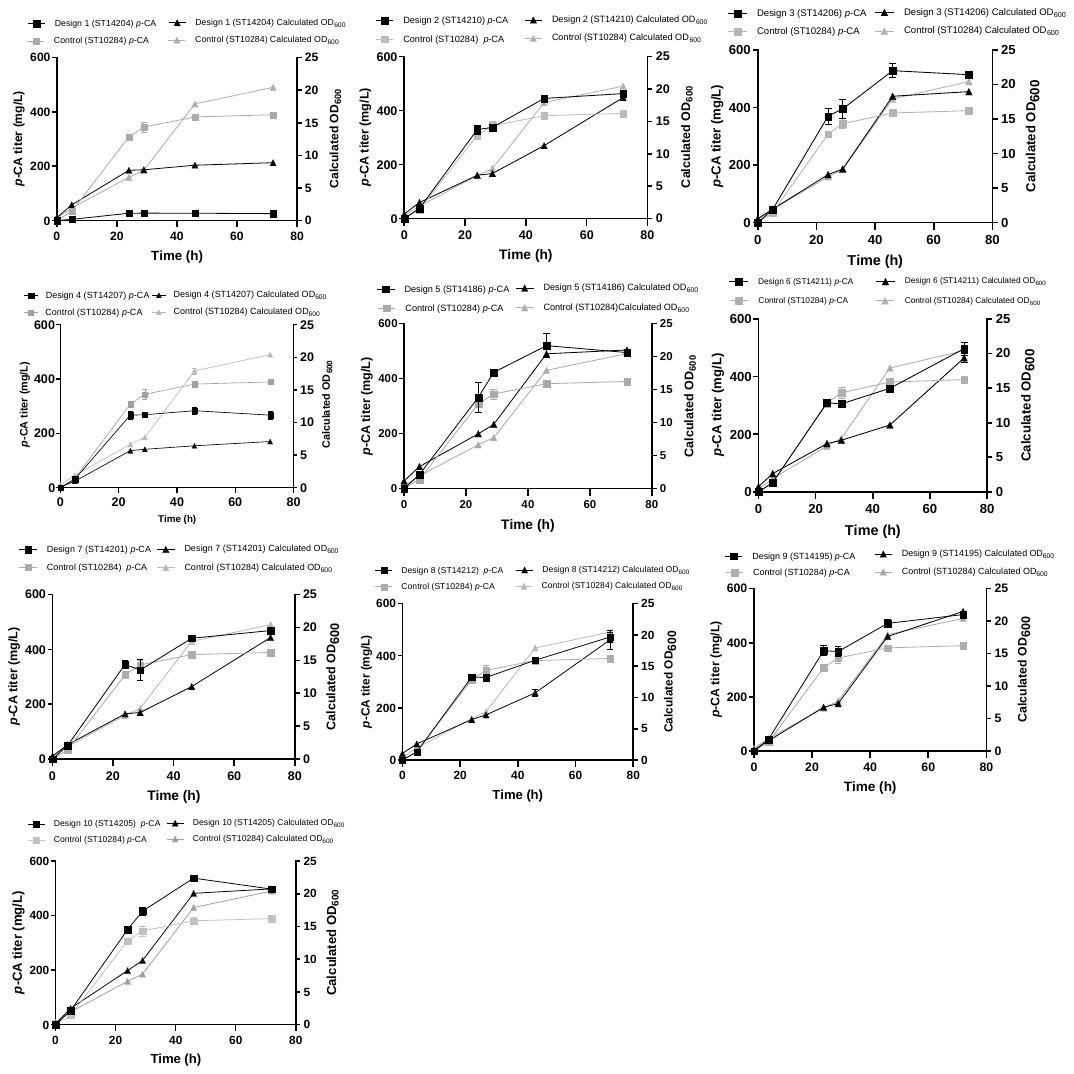
